# Environmental fluctuations reshape an unexpected diversity-disturbance relationship in a microbial community

**DOI:** 10.1101/2020.07.28.225987

**Authors:** Christopher P. Mancuso, Hyunseok Lee, Clare I. Abreu, Jeff Gore, Ahmad S. Khalil

## Abstract

Environmental disturbances have long been theorized to play a significant role in shaping the diversity and composition of ecosystems^1,2^. However, fundamental limitations in our ability to specify the characteristics of a disturbance in the field and laboratory have produced an inconsistent picture of diversity-disturbance relationships (DDRs) that shape the structure of ecosystems^3^. Here, using a recently developed continuous culture system with tunable environmental control^4^, we decomposed a dilution disturbance into intensity and fluctuation components^5,6^, and tested their effects on the diversity of a soil-derived bacterial community across hundreds of replicate cultures. We observed an unexpected U-shaped relationship between community diversity and disturbance intensity in the absence of fluctuations, an observation counter to classical intuition. Adding fluctuations erased the U-shape and increased community diversity across all disturbance intensities. All of these results are well-captured by a Monod consumer resource model, which further reveals how U-shaped DDRs emerge based on a novel “niche flip” mechanism in which competitive outcomes flip and coexistence regimes subsequently collapse at intermediate disturbance levels. Broadly, our results demonstrate how distinct features of an environmental disturbance can interact in complex ways to govern ecosystem assembly and produce all the major classes of DDRs, without invoking other organizational principles. With these findings, we construct a unifying framework that reconciles the disparate DDRs observed in nature, and propose strategies for predictively reshaping the compositional complexity of microbiomes and other ecosystems.

## Introduction

Biodiversity is a cornerstone of ecosystem stability and function^7^. While it is well-appreciated that environmental changes influence species diversity in all ecosystems, the exact nature of this critical relationship is unclear. Without a predictive understanding of how ecosystems respond to perturbations, we are poorly prepared for environmental changes of anthropogenic origin, such as rising global temperatures^8^, and unable to design effective and robust interventions in ecosystems, such as microbiomes of medical or agricultural importance^9,10^. Accordingly, there have been many efforts aimed at understanding the role of environmental disturbances, which bring about mortality of organisms and a reduction of biomass of a community. Various diversity-disturbance (DDR) relationships have been proposed that draw intuition from observations of natural ecosystems. A famous example is the Intermediate Disturbance Hypothesis^1,2^, in which species diversity peaks at intermediate disturbance intensities (**Fig. 1a**). However, DDRs derived from observational studies of disparate ecosystems and disturbance regimes often have inconsistent results^3,11^. Earlier assertions that disturbance weakens or interrupts competition^1,2^ have been refuted by both theory^12,13^ and empirical findings^14^ that harsher environments instead reinforce dependence on limiting factors. Without a framework that directly pairs theory and experiment, it has been difficult to determine the source of disagreement between the many conflicting predictions and observations surrounding DDRs.

**Fig. 1.**
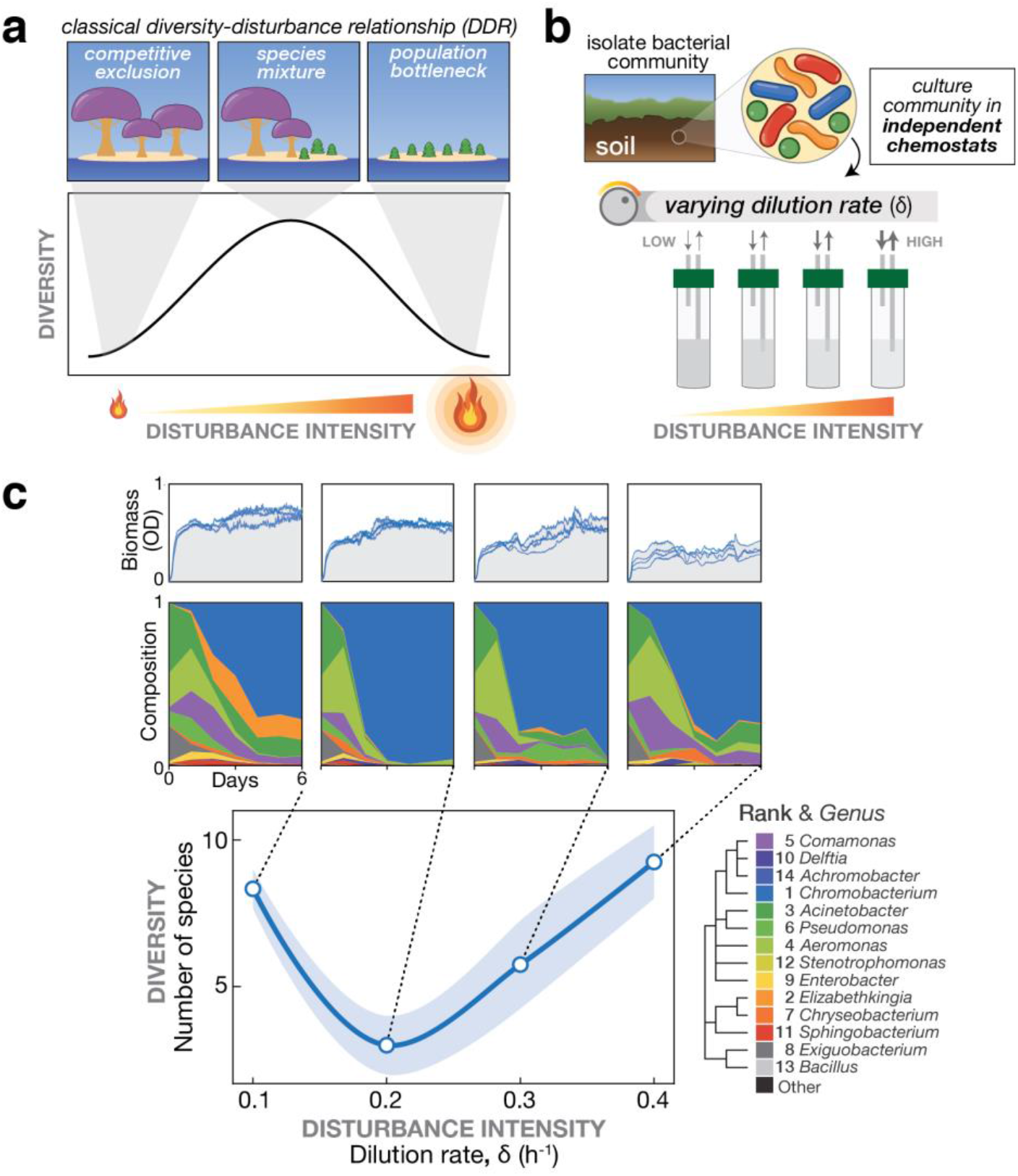
Emergence of a U-shaped diversity-disturbance relationship (DDR) in a microbial community for constantly applied disturbance at different intensities. **(a)** Different DDRs have been proposed based on observations of natural ecosystems, including the Intermediate Disturbance Hypothesis in which diversity peaks at intermediate disturbance levels, see depiction. **(b)** In the laboratory, microbial communities can be cultivated and subjected to varying disturbance intensity levels by tuning the dilution rate in chemostats. **(c)** A bacterial community exhibits a U-shaped diversity dependence on the disturbance intensity. Samples of a soil-derived bacterial community were grown for 6 days in eVOLVER mini-chemostats at four different dilution rates. Top: Optical density over time quantifies biomass for replicate cultures. Middle: Mean relative abundance of bacterial genera from replicate cultures, determined by 16S sequencing. Mean rank abundance is denoted by order, taxonomic similarity is denoted by color. Bottom: Plotting the endpoint number of species (Amplicon Sequence Variants) vs. dilution rate produces a U-shaped curve, rather than a peaked DDR. Shaded window indicates a one standard deviation confidence interval.

Importantly, the impact of a disturbance on an ecosystem depends on the disturbance characteristics. For example, environmental disturbances often introduce fluctuations. The environmental fluctuations associated with a disturbance may in fact stabilize communities by creating temporal niches, similar to seasonal effects^15,16^. Indeed, coexistence can be promoted in a fluctuation-dependent manner due to storage effects (e.g. dormancy in poor conditions) or if species exhibit relative non-linearities in their competitive responses (e.g. differently shaped growth curves)^16,17^. Yet, coexistence might also arise from the overall time-averaged disturbance intensity in a fluctuation-independent manner^12,13^. To determine whether the effects of disturbance on diversity are truly fluctuation-dependent^18^, a disturbance should be decomposed into distinct components of ***intensity*** and ***fluctuation***. Indeed, experimental findings^6,19^ and theory^5^ have suggested that diverse DDRs could arise when considering these factors independently. There is therefore a need for comprehensive, controlled studies which pair theory with experimental methods to produce datasets that can deconvolve the effects of the two key disturbance characteristics: intensity and fluctuation.

Laboratory experiments offer a greater degree of control and throughput compared to field studies, particularly for tractable ecosystem models like microbial communities^20^. Microbes are easily quantified with next-generation sequencing^21-24^, and have been widely used in the laboratory to model community assembly^25-27^, cross-feeding relationships^28^, and succession^29^. Laboratory models have also linked changes in diversity in response to fluctuating nutrient levels^30,31^ and disturbances such as sonication^14^, ultraviolet radiation^32^, osmotic pressures^17^, or toxic compounds^33^. Dilution is perhaps the most common choice for a laboratory disturbance, as it causes species-independent mortality and replenishes the system with fresh nutrients, reminiscent of flow in soil, aquatic, or gut microbiomes. In simple batch culture experiments, where cultures remain undisturbed except for a periodic dilution step, coexistence has been observed at intermediate dilution levels^32,34^, though different DDRs arise under different dilution regimes^6^, suggesting that the dilution parameter space is vastly under-sampled. For more precise tuning of dilution or other parameters, experimentalists have long relied on continuous culture methods^30,31,35^; unfortunately, these systems have traditionally been intractable to large-scale, multidimensional experiments. Recently, we developed eVOLVER, a flexible and automated continuous culture platform that enables independent control over conditions in a large number of mini-bioreactors^4,36^, thus opening up the possibility to explore microbial community dynamics under controlled, multidimensional environmental disturbances. By programming different dilution profiles with eVOLVER, we set out to independently quantify the effects of disturbance intensity and fluctuations on the composition and diversity of microbial communities.

First, we sought to measure microbial diversity at various levels of disturbance intensity in the absence of fluctuations. We cultivated replicate samples of a soil-derived microbiome in separate eVOLVER bioreactor arrays in dilute Nutrient Broth for six days (comprising 20-90 generations), during which continuously diluted cultures approached equilibrium. In a chemostat, the flow of media into the vessel is matched by flow of spent media and cells out of the vessel, so disturbance intensity is directly related to dilution rate (**Fig. 1b**). We thus varied the disturbance intensity by varying the dilution rate across the arrays (see Methods, Supplementary Fxgigs. 9 and 10). We sampled cultures daily and used 16S sequencing to quantify composition and diversity over time. As expected, we observed decreasing biomass of the cultures at increasing dilution rates (**Fig. 1c** and Supplementary Fig. 11). Surprisingly, after quantifying the composition of each culture, we observed a U-shaped diversity-disturbance relationship (**Fig. 1c**), with the number of surviving species at intermediate dilution rate at roughly half of the number at either low or high dilution rate. We were particularly intrigued because a U-shaped DDR does not fit the conventional wisdom^2^. Though U-shaped DDRs are rare in empirical observations^3,11^, the conditions under which we observed it were quite straightforward: constantly applied disturbance. Thus, to better understand our observation, we sought a modeling framework in which a U-shaped DDR could emerge from constantly-applied disturbance, while still capturing other reported DDR shapes.

We started by examining the simplest case giving rise to a U-shaped DDR, a two-species competition where coexistence breaks down at intermediate disturbance levels (**Fig 2a**). To link changes in disturbance intensity to changes in competitive outcomes, we turned to consumer resource models^37,38^. In consumer resource models, species growth rates are a function of resource concentrations. The range of resource concentrations that can support growth of a species can be graphically analyzed on a Tilman diagram^37^ by defining a Zero Net Growth Isocline (ZNGI). As a population consumes resources, the resource concentrations move toward the ZNGI, where growth rate is equal to the mortality rate (i.e. disturbance intensity) and the population is at equilibrium. Accordingly, higher mortality rates will move the ZNGIs to higher resource levels. The shape of these ZNGIs also predict the outcome of competition, as resource consumption by the population can cause equilibrium resource levels to cross the ZNGI of one species, leading to exclusion by the other (**Fig 2b**). At the boundary of these regions, invasion of a one species by the other becomes possible. Coexistence can be achieved when these invasion boundaries are arranged such that either species can invade the other; in this region consumption brings equilibrium resource concentrations towards the intersection of the ZNGIs (**Fig. 2b**). In a competition between two species, a U-shaped DDR can be generated if this coexistence region disappears at intermediate disturbance intensities. We propose that this is possible if the ZNGIs and invasion boundaries flip as disturbance intensity increases, such that at some intermediate intensity the invasion boundaries align and the coexistence region disappears (**Fig. 2c**). We term this behavior “niche-flip”. Under niche-flip, the winner of competition changes as disturbance intensity varies, concomitantly with resource availability.

**Fig. 2.**
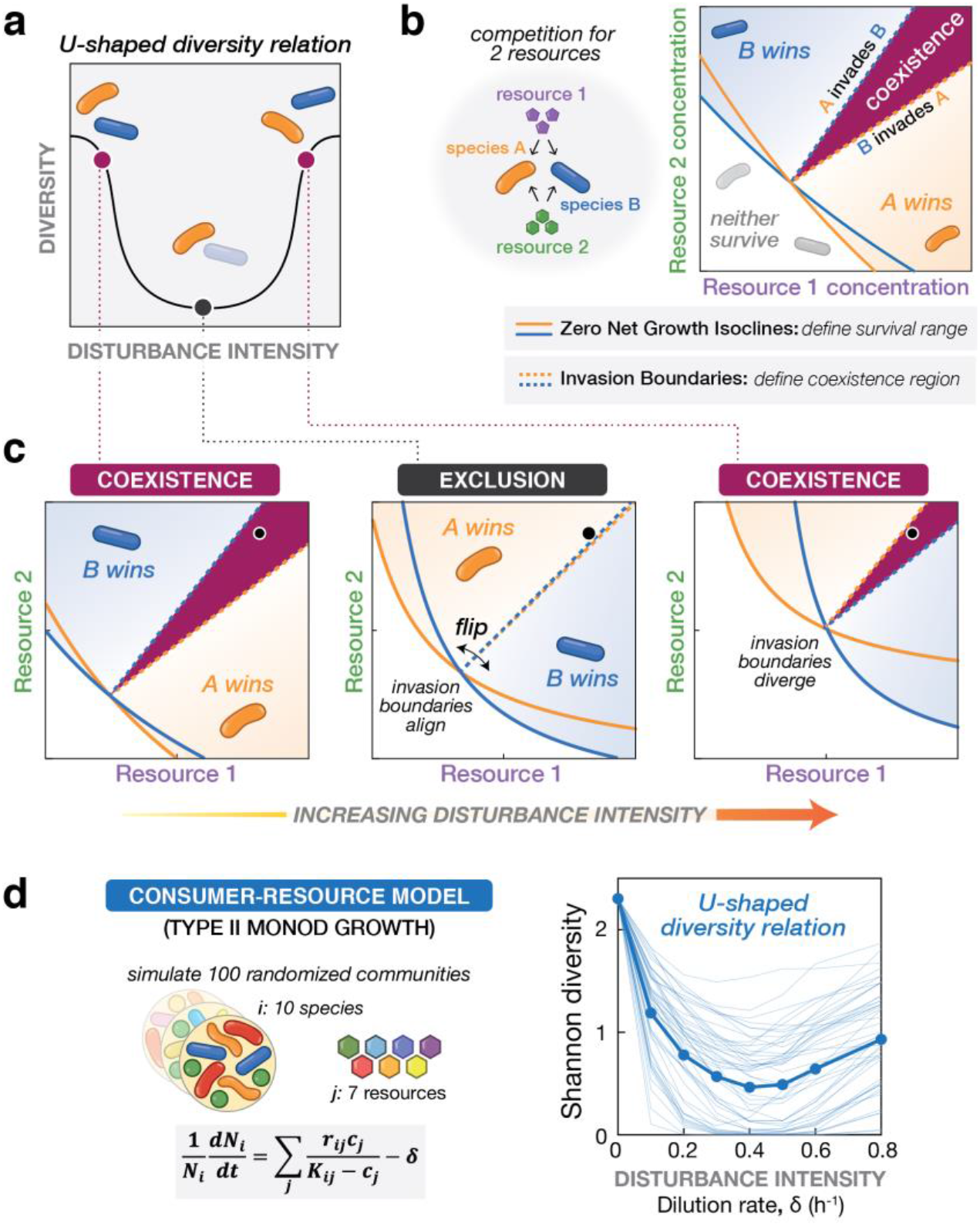
A U-shaped Diversity Disturbance Relationship can emerge from a consumer resource model if species undergo niche-flip as disturbance increases. **(a)** Schematic of a 2-species / 2-resource system showing how a U-shaped DDR could emerge. At low and high disturbance intensities, species must coexist, but at an intermediate level, one species excludes the other. **(b)** The survival of a species in a consumer resource model depends on the supplied resource levels and the mortality rate (i.e. disturbance intensity). A Zero Net Growth Isocline (ZNGI) may be defined for a species at each disturbance intensity, delineating the range of resource supply levels where growth can meet or exceed the mortality rate. Invasion boundaries indicate regions where one species can increase in density in the presence of the other, defining the region of coexistence (maroon). **(c)** In a Type II consumer resource simulation with 2-species / 2-resources, both ZNGIs and resource consumption vectors (defined as consumption by each species at the ZNGI intersection) flip in response to increasing disturbance intensity. At intermediate disturbance, invasion boundaries align and the coexistence region collapses, reducing diversity relative to low/high disturbance intensities. The outcome of competition at the indicated resource supply point (black) is indicated at the top of each plot. **(d)** Left: Monod consumer resource model for growth of species *i* with additive non-linear growth on each resource. Right: Shannon diversity of randomly generated 10-species communities, after six days of simulated growth on seven resources at varying dilution rates. For each model, mean diversity was computed for 100 randomly initialized communities, across each mean dilution rate, 50 of which are shown as individual traces (Methods).

To look for conditions where niche flip emerges, we conducted simulations using Monod growth kinetics^39^, a commonly used model for microbial growth with Type II functional response. Here, growth scales with the concentration of resource, but saturates to a maximal growth rate *r* according to a half-saturation constant *K* (**Fig 2d** and Supplementary Discussion 1). Accordingly, a species with high maximum growth rate *r* may be outcompeted at low resource levels by a species with a low saturation constant *K*, such that the outcome of competition varies depending on nutrient levels (and thus dilution rate δ) (Supplementary Fig. 12). Theoretical analysis of the Monod model with two species and two resources yields ZNGIs and invasion boundaries that undergo niche-flip as dilution rate increases (**Fig 2c**). We simulated sets of 10 species and 7 resources, with per-capita growth rates composed of a sum across nutrient-specific growth rates. Excitingly, we found that the Monod consumer resource model recapitulates the U-shaped diversity dependence on disturbance intensity that we observed in our chemostat experiments (**Fig. 2d**). In systems of larger numbers of species and resources, niche-flip exclusion events between pairs of species can co-occur, yielding U-shaped DDRs (Supplementary Discussion 2). Notably, neither niche flip nor U-shaped DDRs were observed under Lotka-Volterra or linear consumer resource models (Supplementary Discussion 1 and Supplementary Figs. 1 and 2). To confirm that Monod growth kinetics are observed for the species and media conditions of our experiments, we measured growth of isolates in a plate reader. We observed saturating growth rates at higher media compositions and observed variation in *r* and *K* values across resources (Supplementary Fig. 13), consistent with the requirements for niche-flip and fluctuation-dependence (Supplementary Discussion 2). Satisfied that the Monod model could capture the U-shaped behavior of our microbial communities under constant disturbance, we next examined how other DDR shapes could arise under this model.

Both disturbance intensity and fluctuations are hypothesized to play a role in the assembly of ecosystems, but how these two disturbance components interact to reshape DDRs is unclear. Using our modeling framework, we sought to independently vary these two components, simulating a two-dimensional dilution profile. Specifically, we introduced fluctuations into the model by permitting δ to vary with time, compressing disturbance into discrete time windows (**Fig. 3a**); this was done while keeping the time-averaged δ equal, thereby allowing us to vary disturbance intensity and fluctuation independently. The Monod consumer resource simulations predict significantly higher diversity in fluctuating conditions comprised of one or more dilution events per day, with the lowest-frequency (i.e. largest-fluctuation) regime predicted to maintain the most diversity (**Fig. 3d**). This is consistent with intuition that fluctuations introduce temporal structure into environments which may create new niches that promote diversity. Furthermore, the DDR is reshaped entirely – from U-shaped to largely uniform – indicating that community composition in the Monod model is conclusively fluctuation-dependent. Notably, neither Lotka-Volterra nor linear consumer resource models predict differences in the DDR between fluctuation frequencies (Supplementary Figs. 1 and 2). The overlap of DDRs of different frequencies indicates that in these models the relevant metric is the time-averaged overall intensity, rather than the frequency, of disturbance^12,13^.

**Fig. 3.**
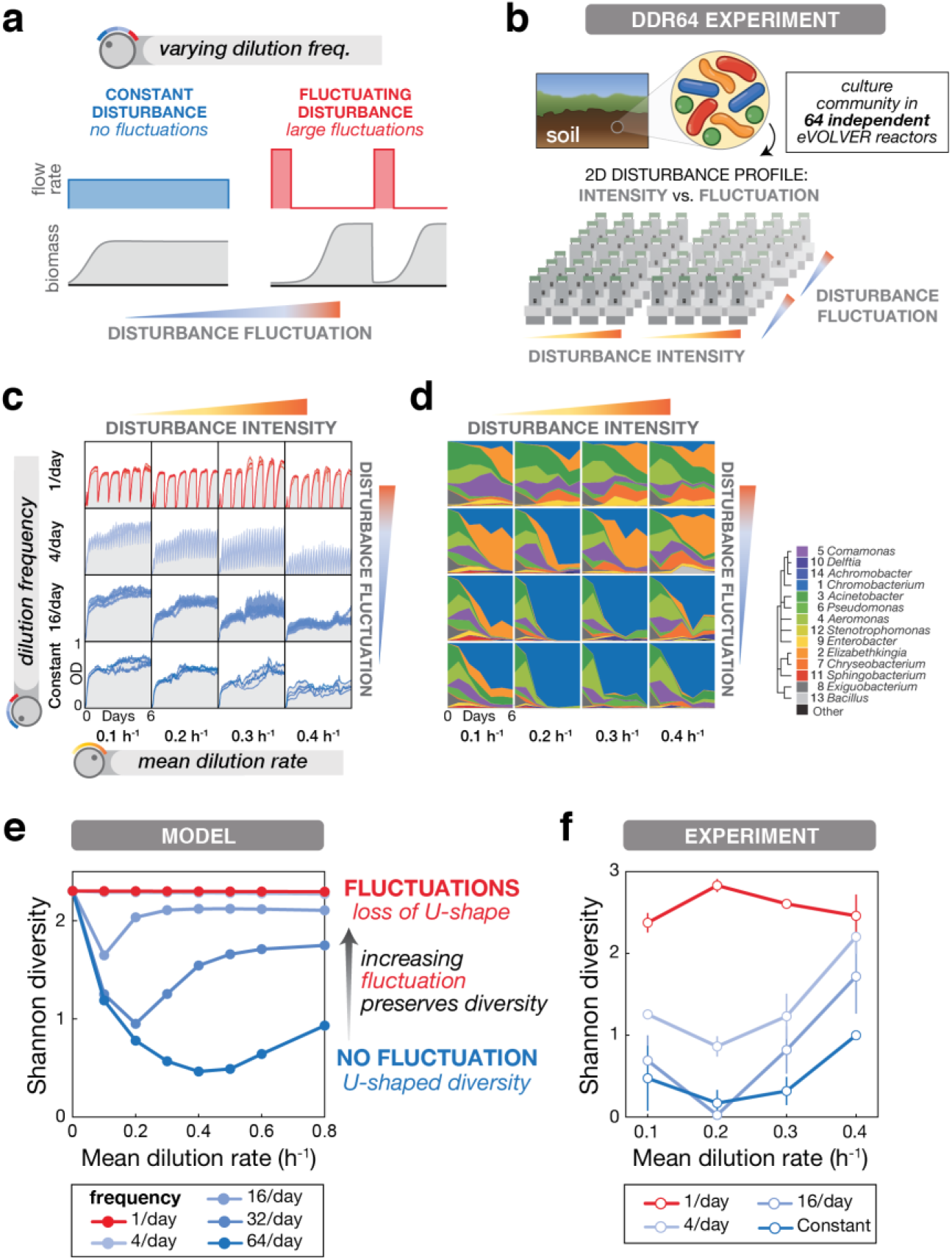
Introducing environmental fluctuations reshapes DDR and increases diversity levels in a microbial community. **(a)** Fluctuations in a disturbance over time cause fluctuations in biomass, and can be varied independently of the disturbance intensity. In continuous culture, fluctuations are achieved by aggregating dilution into discrete events while keeping mean dilution rate constant per day. **(b)** Schematic of the eVOLVER DDR64 experiment in which disturbance components (intensity and fluctuation) are varied independently. Samples of a soil-derived bacterial community were continuously cultured for 6 days across 64 eVOLVER bioreactors with varying mean dilution rate and dilution frequency. **(c)** Optical density traces for culture replicates in each condition show the dependence of biomass on disturbance. **(d)** Mean relative abundance of bacterial genera from replicate cultures, determined by 16S sequencing. Mean rank abundance is denoted by order, taxonomic similarity is denoted by color (see legend). **(e)** Mean Shannon diversity across 100 Monod consumer resource model simulations with varying mean dilution rate and dilution frequency show that the dependence of diversity on disturbance is fluctuation-dependent. **(f)** Mean Shannon diversity of Amplicon Sequence Variants from the DDR64 experiment vs. mean dilution rate and dilution frequencies. As in the simulations, fluctuations increase diversity and eliminate the U-shape. Bars in **e** and **f** denote standard error of the mean.

We returned to experiments to see whether the U-shaped DDR observed in constant dilution chemostats is reshaped by fluctuations, as predicted by the Monod model. To comprehensively test for fluctuation-dependency, we implemented dilution profiles with 1, 4, or 16 fluctuations per day (alongside the constant dilution conditions) at varying mean dilution rates in an eVOLVER experiment comprised of 64 simultaneous cultures (“DDR64 Experiment”) (**Fig. 3b**). As before, we cultivated replicate samples of the soil-derived microbiome in eVOLVER for six days, taking samples every 24 hours to quantify composition. The specific dilution profiles we programmed were reflected in the optical density traces of each culture over time, showing differences between conditions but close agreement between replicates (**Fig. 3c** and Supplementary Fig. 11). Based on 16S sequencing^21-24^, we observed that the genus-level composition of the community varied over time and between conditions (**Fig. 3d** and Supplementary Figs. 14 and 15). Culture compositions diverged from the initial composition, and Principle Coordinate Analysis revealed that constant dilution conditions and 1/day fluctuations diverged from each other, indicating a clear fluctuation-dependent effect, with the spread modulated by mean dilution rate (Supplementary Fig. 15). Notably, despite starting from a diverse community with hundreds of species, we found resulting compositions to be largely similar between replicates (Supplementary Figs. 10 and 14).

We calculated Shannon diversity for each timepoint (Supplementary Fig. 16) and found that endpoint diversity trends across disturbance intensity and fluctuation frequency are qualitatively consistent with the Monod consumer resource model in three ways (**Fig. 3e and f**, and Supplementary Fig. 17). 1) We observed U-shaped diversity curves in regimes of constant disturbance and small frequent disturbances, in both experiment and simulations. 2) Larger fluctuations preserved higher levels of diversity, and 3) larger fluctuations reshaped the diversity-disturbance curves towards a more uniform relationship. Our experimental results were reproducible from frozen inoculum, as confirmed by a 48-vial experiment designed to examine washout at extreme dilution rates (up to 1.5 h^-1^) (Supplementary Discussion 3, and Supplementary Figs. 9, 11, 14, 16, and 17). Though other measurables varied across the disturbance parameter space (e.g. biofilm, DNA content), they do not explain the differences in diversity as clearly as the Monod consumer resource model does (Supplementary Discussion 3, and Supplementary Figs. 18-20). We found it striking that the model captures the features of our results so well while being relatively simple and non-parameterized.

We set out to develop a conceptual framework that not only captures our results, but also provides mechanistic intuition about how the characteristics of disturbance, such as frequency, can produce and reshape DDRs more generally. In this case, we demonstrated how a U-shaped DDR emerging under constantly-applied disturbance was reshaped to a more neutral relationship by the addition of fluctuations. This framework further identified a niche-flip mechanism as a dominant factor in community assembly under constantly applied disturbance. We next wondered whether this simple model could produce other classes of DDRs observed in nature^3^, as a path towards identifying the organizational mechanisms in other ecosystems. To explore a wider range of behaviors, we extended the range of simulations to include extreme disturbance intensities that eventually lead to community extinction, and prevented artificial extinction of all species simultaneously by introducing noise into the normalization of growth rates. As depicted in a contour plot (**Fig 4**), we again found that fluctuations reshaped the DDRs to produce complex diversity landscapes. These simulations revealed that a diverse repertoire of DDRs can emerge from the combination of these two disturbance characteristics (intensity and fluctuations). By exploring subsets of this parameter space, we observed every major class of

DDR: positive, negative, neutral, peaked, and U-shaped (**Fig 4**). By simply examining these two disturbance characteristics in a systematic way, we developed a unifying framework that can reconcile disparate DDR observations without invoking more complex phenomena.

**Fig. 4.**
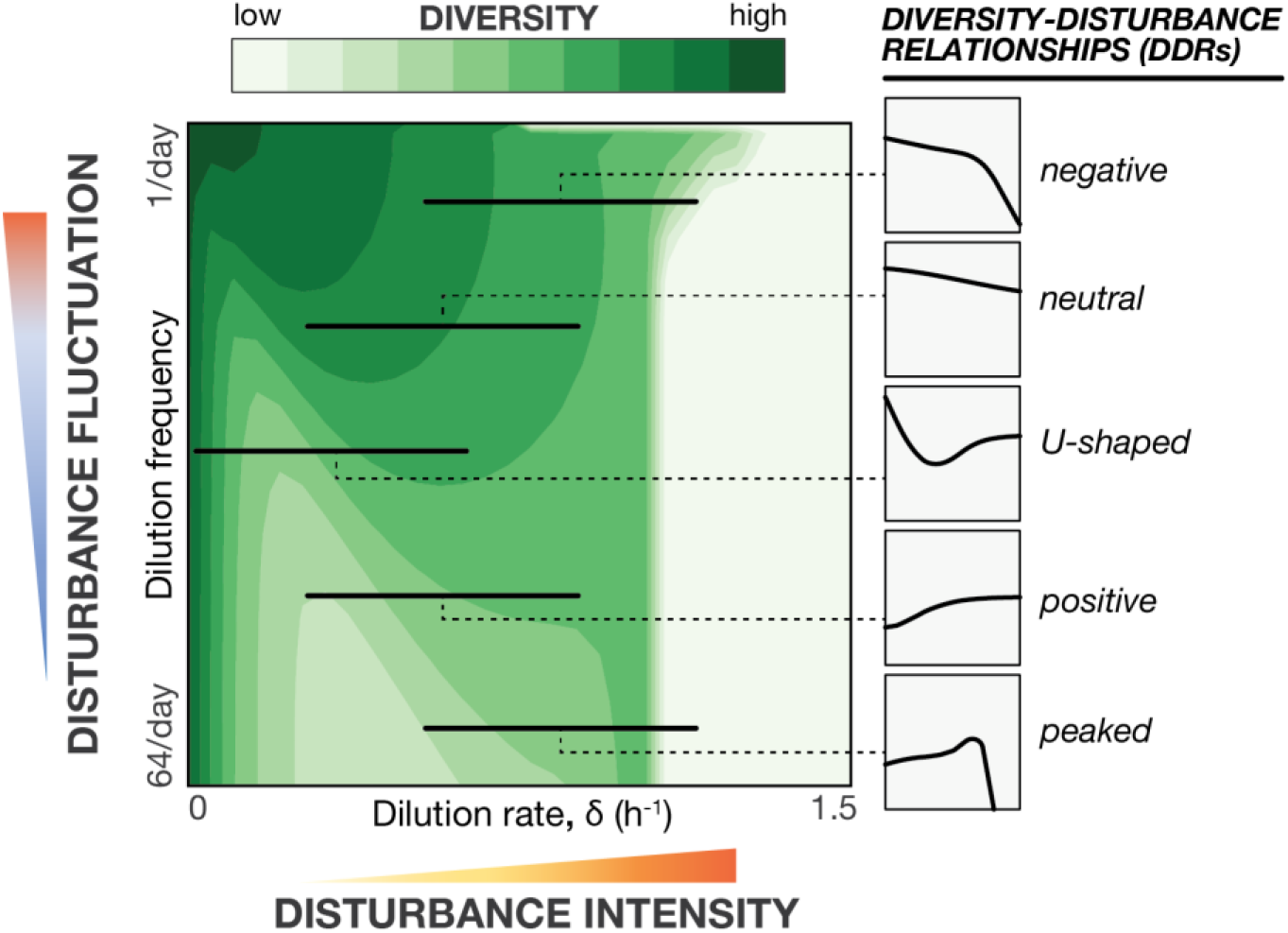
Distinct classes of Diversity-Disturbance Relationships emerge when traversing different regions of disturbance (intensity vs. fluctuation) space. A simple Monod consumer resource model maps a phase diagram for community diversity by varying distinct characteristics of environmental disturbance (intensity and fluctuation). Diversity varies non-linearly with disturbance intensity under different levels of fluctuation for Type II consumer resource models. Contour plot depicts Shannon diversity results for a Monod consumer resource model simulation as an illustrative example. When varying disturbance intensity at a fixed fluctuation level along the indicated arrows, different DDR classes emerge, as shown in smoothed plots on the right.

In this work, advances in automated continuous culture technology and next-generation sequencing enabled laboratory microbial ecology studies to systematically dissect the role of environmental disturbance (intensity and fluctuation) with fine resolution at scale. We found replicable patterns in composition and diversity of a soil-derived microbial community across different disturbance regimes. Notably, we observed an unexpected U-shaped DDR under constant disturbance, and found that adding fluctuations increased community diversity and reshaped the DDR. All of these results are well captured by the Monod consumer resource model, which subsequently led us to describe and propose a novel niche-flip mechanism for structuring these ecosystems. Taken together, these experimental and modeling results 1) provide new insight into how community assembly depends on environmental conditions and 2) demonstrate a role for environmental fluctuations in promoting diversity. Examining the behavior of the Monod model over a broad range of disturbance intensities and fluctuations, we see that diverse classes of DDRs (including increasing, decreasing, and peaked DDRs) could emerge when only subsets of the parameter space are sampled (**Fig 4**). Understanding how disturbance intensity and fluctuations interact is essential for reconciling disparate observations of DDRs under a single unifying framework^3,5,6,11^.

With further study and evaluation of underlying assumptions, the findings of this work may be extended to other systems. Though our experimental results were reproducible (Supplementary Figs. 14, 16 and 17), it remains to be seen whether other species (microbial and macroscopic) and disturbance types (including asymmetric disturbances like toxins or heat shock) behave similarly. The generalizability of the niche-flip mechanism can be evaluated by reexamining the underlying assumptions and formulations of the Monod growth model (Supplementary Discussions 1 and 2). Importantly, we have found that the U-shape is robust to several alternative normalizations and parameter ranges (Supplementary Figs. 3 and 5-8) and non-additive formulations of Monod growth on mixed substrates^40^ (Supplementary Fig. 4). We also found that addition of asymmetric disturbance or introducing migration rates do not qualitatively change our results (Supplementary Fig. 4). Furthermore, the growth saturation feature of the Monod model that enables niche-flip is also characteristic of Type II & III functional responses used in consumer resource models across different scales of ecology. It remains to be seen whether niche-flip mechanisms could arise from non-resource-based models. It is plausible that similar growth tradeoffs arising in response to other disturbance-correlated features could lead to loss of coexistence at intermediate disturbance intensities. Therefore, niche flip could be a more general principle extending beyond relative growth non-linearities explored in this work to systems driven by dynamic abiotic stresses and/or storage effects^41-44^.

Broadly, our results highlight that the structure of ecosystems and their response to perturbation is contextual. We demonstrated that increasing the disturbance intensity can increase, decrease, or have no effect on the diversity of a system. Critically however, we found that under a unifying framework that considers both disturbance intensity and fluctuation these relationships become predictable rather than idiosyncratic^5,6^. Intriguingly, these complex ecosystem assembly rules can emerge from temporal structure alone, without invoking other organizing principles, such as spatial structure^45^ or network structure (e.g. cross-feeding and antagonism)^46,47^. If predictable response to perturbation depends on context, then designing predictable interventions to ecosystems (in medicine, agriculture, and conservation) will require the ability to measure and understand the environmental context. With the staggering amount of compositional data being generated with high throughput sequencing^48^, inference of environmental context and design of robust ecological interventions may not be far off.

## Methods

### Preparing Inoculum

2g of dirt from the Communications Lawn of Boston University (collected on 09/15/2018) was vortexed in 10 mL PBS + 200 ug/mL cycloheximide, then incubated in the dark at room temperature for 48 h. For pre-enrichment, 16 eVOLVER vials were prepared with 25 mL of 0.1X Nutrient Broth (NB) media (0.3 g/L yeast extract + 0.5 g/L peptone (Fisherbrand)) with 200 ug/mL cycloheximide, inoculated with 350 uL of PBS immersion, and grown for 18 h in eVOLVER at 25°C. All 16 pre-enrichment cultures were mixed together to form the experiment inoculum. Aliquots in 15% glycerol were stored frozen at −80°C.

### Running eVOLVER Experiments

eVOLVER lines were sterilized using 10% bleach and ethanol^36,49^, then autoclaved vials were loaded with 23 mL of 0.1X NB. Each vial was inoculated with 1 mL of inoculum, and grown for 5 h at 25°C with stirring prior to the first dilution disturbance. eVOLVER was operated in chemostat mode with 0.5 mL bolus size, with dilutions either evenly distributed over time (constant disturbance) or concentrated in fluctuation periods lasting 15 minutes. For these cultures, the flow rate during a fluctuation *δ_f_*-depended on the number of fluctuations per day *f* and mean dilution rate *δ* according to the following equation: *δ_f_* (24 * *δ*)/(0.25 * *f*) At the end of each experiment, vials were flushed with media, and 10 optical density measurements were taken in eVOLVER to measure the biofilm levels.

Bottles and lines were routinely checked for contamination. This occurred to only a single vial of the experiment, which was excluded from statistical analysis. For the follow-up washout experiment, the glycerol stock inoculum was thawed at room temperature, 1 mL was inoculated into each vial, then the cultures were allowed to grow for 5.7 h prior to initiating disturbances. For the washout experiment, a software bug caused a few incorrectly executed dilution events; these vials were excluded from statistical analysis. Code required to execute these experiments will be available on Github (github.com/khalillab).

### Sampling Cultures

At each timepoint, a 2 mL culture aliquot was removed from each vial with an extended length pipette tip.

For plating, 20 uL of the sample was used for a 10-fold serial dilution series, and 100 uL of diluted cultures at three concentrations were plated on 18 mL Nutrient Broth Agar plates (3 g/L yeast extract, 5 g/L peptone, 15 g/L agar (Fisherbrand)), which were grown at room temperature for 48-60 h, then imaged on an on an Epsom Perfection 550 scanner. Image analysis was performed with the aid of Cellprofiler 3.1.8^50^ and Cellprofiler Analyst 2.2.1 Classifier^51^ tools.

For DNA extraction, the remainder of the sample was pelleted and frozen at −20°C. 60-72 h after freezing, pellets were lysed at 37°C for 1 h in 200 uL of lysozyme buffer (25mM Tris HCl pH 8.0, 2.5mM EDTA, 1% Triton X-100 with 20 mg/mL lysozyme (Fisher), prepared fresh daily). Lysates were processed using DNEasy Blood and Tissue Kit according to manufacturer specifications, eluted into 10 mM Tris buffer, and normalized to 5 ng/uL DNA based on measurements in a Qubit fluorometer.

### Library Preparation and Sequencing

Briefly, we performed amplicon sequencing of the 16S v4 region based on established protocols^21^. Primers prCM543 (TCGTCGGCAGCGTCAGATGTGTATAAGAGAC-AGGTGYCAGCMGCCGCGGTAA) and prCM544 (GTCTCGTGGGCTCGGAGATGTG-TATAAGAGACAGGGACTACNVGGGTWTCTAAT), adapted from EMP515F^52^ and EMP806R^53^ were used to isolate a 290 bp 16S v4 region, using Kapa Hifi ReadyMix polymerase and the following cycling conditions: (i) denaturation: 95°C for 5 min; (ii) amplification (25 cycles): 98°C for 20 s, 55°C for 15 s, 72°C for 1m; (iii) elongation: 72°C for 5 min. For the negative control and biofilm samples, the number of cycles was increased to 35 to amplify from low biomass. Illumina NexteraXT primers (or equivalents) were used to form a final library 427 bp in length, with the following conditions: (i) denaturation: 95°C for 5 min; (ii) amplification (8 cycles): 98°C for 20 s, 55°C for 15 s, 72°C for 1m; (iii) elongation: 72°C for 10 min. DNA was purified with AMPure XP beads or SequalPrep plates, then samples were multiplexed in groups of 192 alongside control samples at a higher fraction, and spiked with PhiX or whole genome DNA libraries to a final concentration of 50% to increase sequence diversity. Library pools were sequenced at the Harvard Biopolymers Facility across five 250 bp paired end MiSeq v2 runs.

### Sequencing Analysis

Samples were demultiplexed using the Illumina BaseSpace demultiplexer analysis tool. All subsequent bioinformatic analysis was performed in QIIME2 v2020.2^22^. Demultiplexed samples were dereplicated using DADA2 sample inference to tabulate Amplicon Sequencing Variants (ASVs)^23^. Next, for qualitative description of composition, taxonomy (to the genus level) was assigned to each feature by alignment to the SILVA 132 database^54^ using the taxa-barplot plugin. For quantitative analysis, samples with technical issues (e.g. contamination, low biomass, poor sequence quality, etc.) were removed and the remaining 698 samples were rarefied to 6840 reads. The fragment-insertion plugin was used to generate a rooted phylogenetic tree using the SEPP algorithm^24^. The diversity plugin was used to calculate Shannon diversity, ASV richness, and weighted UniFrac distance^55^, which was used to perform Principle Coordinate Analysis (PCoA).

## Acknowledgments

We thank members of the Gore and Khalil groups for helpful discussions. We thank S. Boswell for sequencing advice, M. Springer and J. Galagan for reagents and equipment used in library preparation, and the Harvard Biopolymers Facility for their services.

## Author contributions

C.P.M., J.G. and A.S.K. conceived the study. C.P.M. performed all experiments and sequencing analysis. H.L. and C.I.A. performed theoretical modeling. All authors analyzed results. J.G. and A.S.K. oversaw the study. All authors wrote the manuscript.

## Data and materials availability

All sequencing data is being deposited in the Sequence Read Archive (SRA) and will be accessible with a BioProject accession code (Submission number: SUB7786275). Agar plate images are being deposited in a public image repository. Computer code used to run eVOLVER experiments and for theoretical modeling is available at github.com/khalillab. All other datasets required to produce the results in the current study are included as supplemental data.

## Materials and Correspondence

correspondence and material requests may be addressed to A.S.K.

## Supplementary Information

Supplementary Methods

Supplementary Discussions

Supplementary Figures 1-20

