## Supplemental Text and Figures for "Environmental fluctuations reshape an unexpected diversity-disturbance relationship in a microbial community"

### Supplementary Methods

### Supplementary Discussions

**Supp. Discussion 1:** Comparison of simulation results under different microbial competition models . 3
**Supp. Discussion 2:** Theoretical analysis of the niche-flip mechanism underlying U-shaped Diversity Disturbance Relationships.......... 12
**Supp. Discussion 3:** Additional Experimental Results .......... 19

### Supplementary Figures

**Figure 1:** Lotka-Volterra models do not exhibit strong relationship between diversity and fluctuation 4
**Figure 2:** Linear Consumer Resource models do not exhibit relationship between diversity and fluctuation .......... 6
**Figure 3:** Comparison between Monod consumer resource models with variant parameter choices reveal those that retain U-shape .......... 8
**Figure 4:** Comparison between Monod consumer resource models with variant assumptions can retain U-shape .......... 11
**Figure 5:** Generating species with sorted $\boldsymbol{r}$ and $\boldsymbol{K}$ parameters prevents intersection of growth curves 15
**Figure 6:** Communities with sorted $\boldsymbol{K}$ per resource still exhibit U-shaped DDR. .......... 16
**Figure 7:** Niche flip occurs in a community with sorted $\boldsymbol{K}$ parameters at high dilution rate .......... 16
**Figure 8:** Communities with constant $\boldsymbol{K}$ across all resources do not exhibit U-shaped DDR .......... 18
**Figure 9:** Timeline of DDR64 and DDR Washout Experiments .......... 22
**Figure 10:** Inoculum composition determined by 16S sequencing .......... 23
**Figure 11:** Optical density over time for DDR64 and DDR Washout experiments .......... 24
**Figure 12:** Navigating between Monod growth curves and Tilman diagrams .......... 25
**Figure 13:** Measured growth of isolates in Nutrient Broth indicate variation in r and K parameters across different medias .......... 26
**Figure 14:** Genus-level composition over time in the DDR64 and DDR Washout experiments .......... 27
**Figure 15:** Principle Coordinate Analysis shows varying composition in DDR64 Experiment .......... 28
**Figure 16:** Diversity changes over time in DDR64 and DDR Washout experiments .......... 29
**Figure 17:** Different diversity metrics from DDR64 and DDR Washout experiments show U-shaped DDRs .......... 30
**Figure 18:** Heatmap of biofilm levels at endpoint .......... 31
**Figure 19:** Diversity does not correlate with potential confounding factors .......... 32
**Figure 20:** Correlation between population size estimates vary according to condition and biofilm accumulation .......... 33

### Supplementary Methods

Isolating Strains

Colonies from imaging plates for the inoculum and endpoint timepoints of the DDR64 and follow-up washout experiments were re-struck on NB agar plates, grown overnight in 0.1X NB media, and stored frozen at -80˚C in 15% glycerol. Primers prCM215 (CCATTGTAGCACGTG-TGTAGCC) and prCM216 (ACTCCTACGGGAGGCAGC) were used to amplify the v3-v7 16S region for Sanger sequencing and identification.

Growth Characterization of Isolates

Several strains with different taxonomic background were selected from the collection of isolates for further study. For measurement of *r*/*K* values, 9 strains were struck onto NB plates, and single colonies were grown to stationary phase overnight at 30˚C, then diluted to an OD600 of 0.001 in triplicate 200 uL cultures in dilute NB media (ranging from 0.1X to <0.0016X, plus a water-only negative control) in 96-well microplates grown in a Tecan spectrophotometer for 36 h at 30˚C, shaking, with lid on. Growth rates were fit to log transformed OD600 data, Lineweaver-Burke plots were constructed across media concentration, and Monod curves were fit to each species. Three strains that could grow in M9 defined minimal media were selected for characterization on different limiting resources. Strains were struck onto NB plates, and single colonies were grown to stationary phase overnight at 30˚C in carbon-limited M9 supplemented with aspartate, glutamate, or proline, then diluted and grown in microplates as above, with amino acid concentrations ranging from 10 mM to <0.16 mM, plus a no-carbon control. Due to limited growth on these media, optical densities at sub-saturating resource concentrations were below the limit of detection, so only maximum growth rate at 10 mM amino acid was reported.

### Supplementary Discussion

#### Discussion 1. Comparison of simulation results under different microbial competition models

Here we describe the methods, equations, and parameter choices used in our simulation analyses. We explored two general types of microbial competition model: Lotka-Volterra and consumer resource. Within consumer resource models, we explored linear growth models and Monod growth models. We ran further simulations with a number of variations on the Monod growth consumer resource model as this model best fit our observations and proved to be most responsive to the disturbance intensity and fluctuation parameters we investigated.

Lotka-Volterra Simulations

We simulated 10-species competitions with the Lotka-Volterra model:

$$\frac{1}{N_i}\frac{dN_i}{dt} = r_i\left(1 - \sum_{k=1}^{10} \alpha_{ik} N_j\right) \qquad Eqn. 1$$

$N_i$ represents the abundance of species $i$ modified by its carrying capacity (we rescale the carrying capacities of all species to one), $r_i$ represents its maximal growth rate, and $\alpha_{ik}$ represents inhibition of species $i$ by species $k$. The model is parameterized such that $\alpha_{ii} = 1$.

Simulations mimicked experimental conditions as much as possible. Beginning with equal abundances of all 10 species, we integrated equations for a six-day period using the function ode45 in MATLAB. Species abundances were diluted during equally spaced 15-minute intervals by integrating a version of Eqn. 1 modified to include dilution:

$$\frac{1}{N_i}\frac{dN_i}{dt} = r_i\left(1 - \sum_{k=1}^{10} \alpha_{ik} N_k\right) - \delta \qquad Eqn. 2$$

$\delta$, the death or dilution rate, was calculated by distributing the mean dilution rate $D$ (per hour) over equally spaced intervals of 15 minutes (per 1/4 hour) at a frequency $f$ (per 24 hours):

$$\delta = \frac{1}{\left(\frac{1}{4}\right) hr} * \frac{D\left(\frac{1}{hr}\right)}{f\left(\frac{1}{24hr}\right)} = 96 * \frac{D}{f}\left(\frac{1}{hr}\right) \qquad Eqn. 3$$

Frequency $f$ of dilution ranged from 1 to 64 per day, and mean dilution rate $D$ ranged from 0.1 to 0.8 per hour. Maximal growth rates $r_i$ were randomly sampled from a normal distribution with mean 1 and standard deviation 0.1. This range was selected to roughly match measured growth rates of isolates on 0.1X Nutrient Broth (fig. S17). Competition coefficients $\alpha_{ik}$ were randomly sampled from a lognormal distribution with parameters $\mu = -0.7$ and $\sigma = 0.2$, which are the mean and standard deviation of the associated normal distribution. The mean of the lognormal distribution is $\exp\left(\mu + \frac{\sigma^2}{2}\right) = 0.51$ and the standard deviation is $\exp(2\mu) +$

$\sigma^2(\exp(\sigma^2) - 1) = 0.1$. The competition coefficients were selected to match the diversity of the resource-explicit simulation results at the zero-dilution condition (see below). Other ranges of $r_i$ and $\alpha_{ik}$ did not alter the DDR shape (Supplementary Fig. 1).

We simulated 100 competitions with randomly drawn parameters, across all dilution-frequency combinations. All 10 species began at equal abundances, and we used their final relative abundances $p_i$ to calculate the Shannon diversity index $\rho$ of each outcome:

$$\rho = \sum_{i=1}^{10} -p_i \ln p_i \qquad Eqn. 4$$

In order to be counted, the abundance of a species $p_i$ had to exceed a threshold of 0.0001. Finally, we took the average of the 100 values of $\rho$ for each dilution-frequency combination to obtain average diversity.

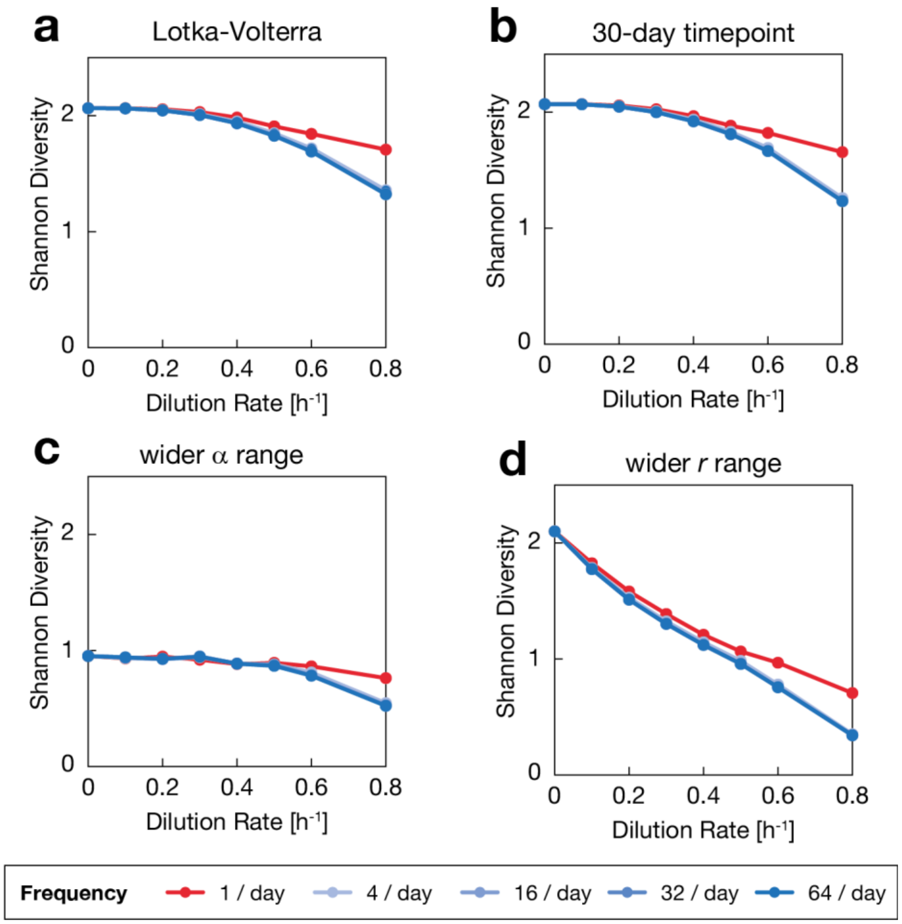


**Supplementary Figure 1. Lotka-Volterra models do not exhibit strong relationship between diversity and fluctuation.** Shannon diversity was plotted vs. disturbance intensity (mean dilution rate), taking the mean across 100 communities after 6 days of simulated growth.

Fluctuations are classified by frequency of dilution disturbance, indicated by color. **(a)** Shannon diversity for Lotka-Volterra model shows little relationship between diversity and fluctuations. **(b)** Longer simulation lengths do not change this relationship. **(c)** Simulations were run using growth rate parameters with a wider uniform range ($\boldsymbol{r}$ = 0.1-1) and decreased mean. Though mean diversity dropped, no qualitative change in DDR shape was observed. **(D)** Simulations were run using $\boldsymbol{\alpha}$ interaction parameters with a wider uniform range ($\boldsymbol{\alpha}$ = 0.5-1.5) and increased mean. The DDR was only minorly affected, retaining a decreasing relationship between diversity and disturbance intensity.

Linear Consumer Resource Simulations

We simulated 10-species, seven-resource competitions with a linear resource concentration model:

$$\frac{1}{N_i}\frac{dN_i}{dt} = \sum_{j=1}^{7} r_{ij}c_j \qquad Eqn. 5$$

$N_i$ represents concentration of species $i$, $r_{ij}$ its growth rate per unit resource on resource $j$, and $c_j$ the concentration of resource $j$. Units of species and resources per volume are the same, because we assume that one unit of resource is fully converted into one unit of biomass; this assumption is equivalent to assuming the biomass yield is equal to one for all species, and therefore we do not include a yield parameter.

Similar to the Lotka-Volterra simulations, we integrated the dilution-modified equations over the same time period and range of frequencies and dilutions:

$$\frac{1}{N_i}\frac{dN_i}{dt} = \sum_{j=1}^{7} r_{ij}c_j - \delta \qquad Eqn. 6$$

$\delta$, as defined by Eqn. 3, is not only the dilution rate of cells, but also the influx of fresh resources at source concentration $c_{jo}$:

$$\frac{dc_j}{dt} = \delta\left(c_{jo} - c_j\right) - \sum_{i=1}^{10} N_i r_{ij} c_j \qquad Eqn. 7$$

Simulations began with equal abundances of all species and resources ($c_{jo} = 1$), except for one resource, which had a slightly different supply concentration ($c_{jo}$ =1.2), in order to move the resource supply point away from a unique central position. Growth rates per unit resource $r_{ij}$ were randomly sampled from a normal distribution with mean 1 and standard deviation 0.1, and then normalized by dividing by the sum of growth rates for each species across all resources, $\sum_j^7 r_{ij}$. Other parameter ranges did not alter the DDR shape (Supplementary Fig. 2).

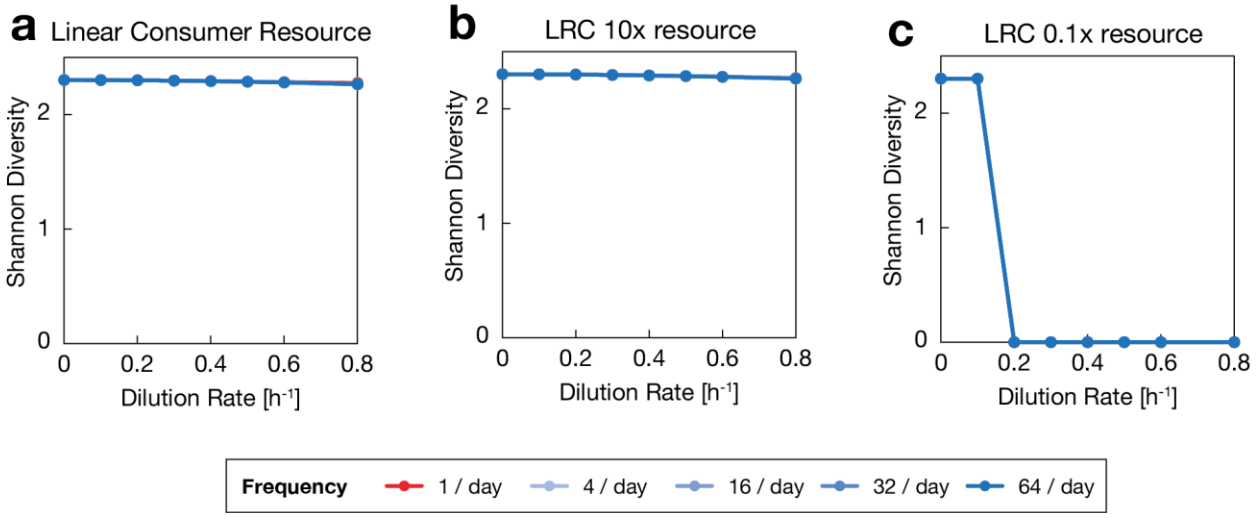


**Supplementary Figure 2. Linear Consumer Resource models do not exhibit relationship between diversity and fluctuation.** Shannon diversity was plotted vs. disturbance intensity (mean dilution rate), taking the mean across 100 communities after 6 days of simulated growth. Fluctuations are classified by frequency of dilution disturbance, indicated by color. **(a)** Shannon diversity for linear consumer resource (LRC) model shows no relationship between diversity and fluctuations. **(b)** Simulations were run with increased resource supply concentrations ($\mathbf{10} \times \boldsymbol{c_{jo}}$), which did not affect the DDR. **(c)** Simulations were run with decreased resource supply concentrations ($\mathbf{0.1} \times \boldsymbol{c_{jo}}$). This reduced growth rates to a level below the dilution rate, leading to washout and a corresponding Shannon diversity of zero.

<u>Monod Consumer Resource Simulations</u>

We simulated 10-species, 7-resource competitions with a Monod resource model:

$$\frac{1}{N_i}\frac{dN_i}{dt} = \sum_{j=1}^{7} \frac{r_{ij}c_j}{K_{ij} + c_j} \qquad Eqn. 8$$

The Monod constant, $K_{ij}$, is the concentration of resource $j$ at which species $i$ reaches its half-maximal growth rate on that resource. Similar to the linear resource concentration model (Eqns. 6-7), the model can be modified to include dilution:

$$\frac{1}{N_i}\frac{dN_i}{dt} = \sum_{j=1}^{7} \frac{r_{ij}c_j}{K_{ij} + c_j} - \delta \qquad Eqn. 9$$

$$\frac{dc_j}{dt} = \delta(c_{jo} - c_j) - \sum_{i=1}^{10} \frac{N_i r_{ij} c_j}{K_{ij} + c_j} \qquad Eqn. 10$$

We performed simulations using the same procedure described above. The Monod constants $K_{ij}$ were randomly sampled from a uniform distribution: [0.001, 0.01]. The width of the selected range is consistent with the range of Monod constants measured on Nutrient Broth (Supplementary Fig. 13), and similar DDR shapes are generated with alternative parameter choices (Supplementary Fig. 3). Maximal growth rates $r_{ij}$ were sampled and normalized as described above. To check whether the U-shaped DDR was preserved under variations of the Monod model, we performed other simulations.

First, we asked whether larger relative fitness differences between species would affect our results or yield different DDRs. In simulations with dilution rates extending out to 1.5 $h^{-1}$, larger than the maximal growth rate of species, we observed unrealistic behavior wherein all species went extinct simultaneously upon reaching the maximum growth rate (normalized for all species). We added Gaussian noise to maximal growth rates $r_{ij}$ with standard deviation of 0.02, which created small differences in relative fitness between species. With this change we observed a more gradual decrease in the diversity as species are lost as dilution rates exceed each species maximal growth rate (**Fig 4**). For larger relative differences, we removed normalization between species, and instead normalized maximal growth rates pre resource by dividing by the number of resources, rather than the sum of growth rates across all resources. In these simulations, the relative differences between species can be quite large. The resulting diversity was even more sensitive to disturbance intensity, causing diversity to decrease steadily with mean dilution rate, but the U shape was preserved (Supplementary Fig. 3e).

Diverging from the conditions of the experiment, we searched for alternative parameter ranges that could qualitatively alter the DDR. We found the model to be robust to increased simulation length (fig. S18d) and increased resource supply (fig. 18b). However, when reducing each of the resource supply concentrations, $c_{jo}$, by a divisive factor of 10, the U-shape was almost completely eliminated (except for a small effect at the highest dilution frequency, fig. S18c). This modification is equivalent to increasing Monod constants, $K$, by a multiplicative factor of 10, placing them closer to the range of resource supply concentration, which does not reflect our measurements (Supplementary Fig. 13) and is less biologically relevant. This extreme parameter range essentially changes the model choice, moving the growth dynamics closer to those of the linear consumer resource model.

Next, we made modifications to parameter choices under the summed Monod model that change the underlying assumptions to see if any would eliminate the U shape. First, we included the possibility of "specialist" species by widening the maximal growth rate sampling range. We randomly sampled $r$ from a uniform distribution (0.1,1), before normalizing by dividing by the sum of growth rates for each species across all resources, $\sum_j^7 r_{ij}$, as before. The U shape was preserved (Supplementary Fig. 3f). Next, we eliminated $r/K$ tradeoffs on all resources by sampling $r$ and $K$ as before, and then sorting them such that the species with the highest value of $r$ also had the lowest value of $K$, and the next-highest and next-lowest, and so on, for each resource. The U shape was preserved (Supplementary Fig. 3g). Finally, we sampled $K$ only once per resource, keeping it constant for all species on each resource. This modification eliminated the U shape (Supplementary Fig. 3h). For a mathematical discussion of the importance of these parameter choices, see Supplementary Discussion 2.

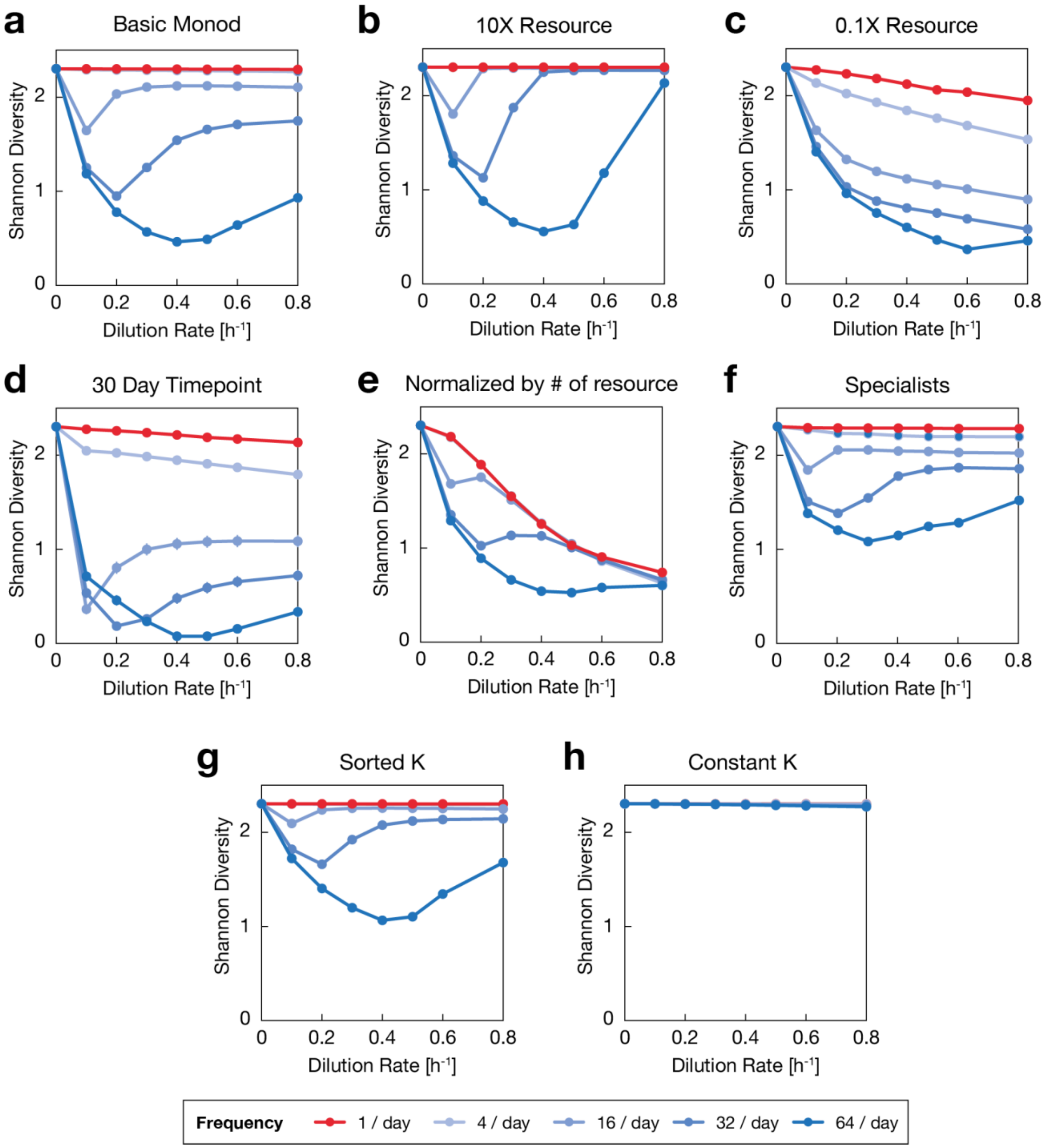


**Supplementary Figure 3. Comparison between Monod consumer resource models with variant parameter choices reveal those that retain U-shape.** Each plot displays Shannon diversity after 6 days of simulated growth, mean across 100 communities for varying mean dilution rate and dilution frequency (see legend). Variations on the basic Monod model are discussed in Materials and Methods. **(a)** Basic Monod Growth model, reproduced from Fig. 3e. **(b)** Increasing resource supply by a factor of 10 does not qualitatively change the DDR. **(c)**

Decreasing resource supply by a factor of 10 strongly affects the DDR by moving the competition out of the saturating nutrient range (set by $\boldsymbol{K}$), such that the Monod model approaches a linear consumer resource model. **(d)** Carrying out the basic Monod growth model from **a** over longer simulation lengths retains the qualitative DDR shape. **(e)** Randomized growth rates on each resource were normalized by the number of resources rather than the species summed growth rates, which permitted wide variance in overall species growth rates, but retained some U-shape. **(f)** Simulations of communities of 10 species of specialists that consumed only 3 out of 7 resources exhibited higher diversity, but retained U-shaped DDRs. **(g)** Sorted $\boldsymbol{K}$ simulations, reproduced from Supplementary Figure 6. **(h)** Constant $\boldsymbol{K}$ simulations, reproduced from Supplementary Figure 8.

Finally, we sought to determine whether changing the functional form of the summed Monod model would affect the U shape. First, we employed a non-additive formulation of Monod growth on mixed substrates[40]. In this formulation, the per-capita growth rate on multiple resources is a saturating function of Monod growth on individual resources, rather than a simple sum (as in Eqn. 10 above):

$$\frac{1}{N_i}\frac{dN_i}{dt} = \frac{\lambda_c \sum_{j=1}^{7} \frac{\phi_{ij}}{\lambda_c - \phi_{ij}}}{1 + \sum_{j=1}^{7} \frac{\phi_{ij}}{\lambda_c - \phi_{ij}}} \qquad Eqn. 11$$

Here, $\phi_{ij} = \frac{r_{ij} c_j}{K_{ij} + c_j}$, which is the Monod growth of species $i$ on resource $j$. The other parameter, $\lambda_c$, is the horizontal intercept in a plot of a species' catabolic enzyme expression as a function of its growth rate. For simplicity, we assumed $\lambda_c$ to be equal for all species. Because the species consume resources non-additively in this model, the resources are also depleted non-additively:

$$\frac{dc_j}{dt} = -\sum_{i=1}^{10} \frac{N_i \phi_{ij}(\lambda_c - \phi_{ij})}{\lambda_c - \frac{1}{N_i}\frac{dN_i}{dt}} \qquad Eqn. 12$$

We can write the dilution-modified model as follows (substituting $\lambda_c \gamma_i$ for per-capita growth of species $i$ in the absence of dilution):

$$\frac{1}{N_i}\frac{dN_i}{dt} = \lambda_c \gamma_i - \delta \qquad Eqn. 13$$

$$\frac{dc_j}{dt} = \delta(c_{jo} - c_j) - \sum_{i=1}^{10} \frac{N_i \phi_{ij}(\lambda_c - \phi_{ij})}{\lambda_c - \lambda_c \gamma_i} \qquad Eqn. 14$$

Simulating this model with values of $\lambda_c$ ranging between 0.3 and 30 yielded DDRs similar to those of the summed Monod model, where the U shape was preserved (Supplementary Figure 4a).

Next, we incorporated the possibility of asymmetric death rates by introducing a species-specific death rate in addition to a uniform dilution rate:

$$\frac{1}{N_i}\frac{dN_i}{dt} = \sum_{j=1}^{7} \frac{r_{ij}c_j}{K_{ij} + c_j} - \delta - \delta_i \qquad Eqn. 15$$

The species-dependent death rate $\delta_i$, randomly sampled from a uniform distribution of [-0.1, 0.1], introduced asymmetry to mortality of each species in the community. Species with high mortality were severely penalized compared to species with low mortality, and the species pool was effectively reduced to only species with low mortality. This modification led to a lower maximum diversity of the system, but U-shaped DDR was maintained (Supplementary Fig. 4b).

Finally, we incorporated the possibility of migration by introducing a species-specific migration from a common source:

$$\frac{1}{N_i}\frac{dN_i}{dt} = \sum_{j=1}^{7} \frac{r_{ij}c_j}{K_{ij} + c_j} - \delta + \frac{m_i}{N_i} \qquad Eqn. 16$$

The migration rate $m_i$, set as 0.001 for all species, is applied continuously regardless of the fluctuation in disturbance, modeling a constant population influx from a highly diverse common source. This uniform migration boosted the diversity overall, yet the U-shaped DDR was preserved (Supplementary Fig. 4c).

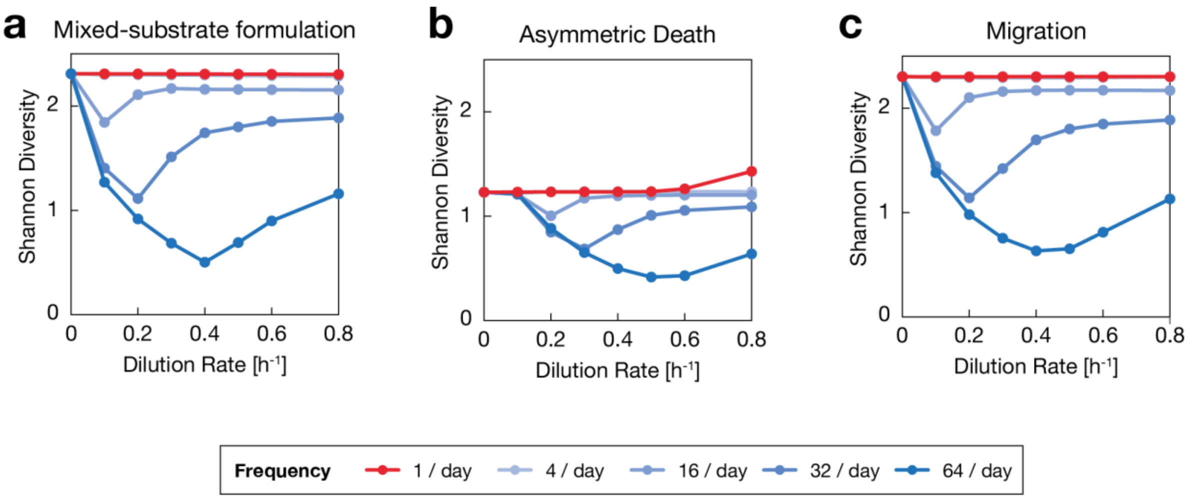


**Supplementary Figure 4. Comparison between Monod consumer resource models with variant assumptions can retain U-shape.** Each plot displays Shannon diversity after 6 days of simulated growth, mean across 100 communities for varying mean dilution rate and dilution frequency (see legend). Variations on the basic Monod model are discussed in Materials and Methods. **(a)** Growth rates across multiple resources were normalized using an alternative mixed-substrate utilization model, retaining the DDR shapes. **(b)** Rather than having equal mortality rates across species, each species was assigned an independent mortality rate $\boldsymbol{\delta_i}$ in addition to the universal dilution rate $\boldsymbol{\delta}$. Diversity is reduced as species with higher mortality rates are penalized, but this does not qualitatively change the DDRs observed. **(c)** To account for migration, each species was assigned a migration rate from a universal source community, independent of the dilution rate. Though species are constantly reintroduced into the community, if the migration rates are low compared to growth and dilution rates, then there are no major qualitative changes to the observed DDRs.

**Supplementary Discussion 2: Theoretical analysis of the niche-flip mechanism underlying U-shaped Diversity Disturbance Relationships**

As discussed in the main text, the U-shaped diversity disturbance relationship we observed under constantly applied disturbance has not been extensively explored in the literature. In a theoretical study based on this observation, we describe how a mechanism we term 'niche-flip' can lead to a U-shaped diversity-disturbance relationship. To introduce niche flip, we focus on a two-species, two-resource community. In this supplementary section, we will mathematically show how *r*/*K* tradeoffs can lead to niche flip. Without losing generality, these conditions can be summarized qualitatively:

- At low disturbance intensities, species 1 drives species 2 extinct in resource 1 and species 2 drives species 1 extinct in resource 2. Species 1 and 2 can coexist in the presence of both resource 1 and 2.
- At high disturbance intensities, species 2 drives species 1 extinct in resource 1 and species 1 drives species 2 extinct in resource 2. Species 1 and 2 can coexist in the presence of both resource 1 and 2.

We assume that the growth rate of each species is represented by a summed Monod model:

$$\frac{1}{N_i}\frac{dN_i}{dt} = \sum_j r_{ij}\,\frac{c_j}{K_{ij} + c_j} - \delta \qquad Eqn. 17$$

$N_i$ is the population of species $i$, $c_j$ is the concentration of resource $j$, $r_{ij}$ is the maximum growth rate of species $i$ on resource $j$, $K_{ij}$ is the Monod constant of species $i$ on resource $j$, and $\delta$ is a global death rate added to the system, such as the dilution rate in our experiments. The magnitude of $\delta$ is the disturbance intensity in the system.

We consider chemostat-like continuous dynamics, where resources are supplied at the same rate at which they are diluted, and they are consumed due to the species' growth:

$$\frac{dc_j}{dt} = -\sum_i a_{ij} r_{ij} \frac{c_j}{K_{ij} + c_j} + \delta\left(c_{j0} - c_j\right) \qquad Eqn. 18$$

$a_{ij}$ is the inverse yield of resource $j$ for species $i$ (i.e. how much of resource $j$ is needed for a unit growth in population $i$), and $c_{j0}$ is the fresh media concentration of supplied resource $j$. For simplicity, we will assume $a_{ij} = 1$ throughout our analysis.

We note that the competition result under a single supplied resource is determined by the equilibrium resource concentration $c_{ij}^*$ when species $i$ survives entirely on resource $j$. $c_{ij}^*$ is then a solution of:

$$r_{ij}\frac{c_{ij}^*}{K_{ij} + c_{ij}^*} = \delta \qquad Eqn. 19$$

Then species 1 drives species 2 extinct in resource 1 if and only if:

$$c_{11}^* < c_{21}^* \qquad Eqn. 20$$

Let us consider two disturbance regimes separately, and describe the necessary conditions for coexistence in the presence of both resources in each case:

***Condition 1***: Low dilution regime: $\delta \ll r_{ij}$

In this case, the resource concentrations at equilibrium are much smaller than the Monod constant: $c_j^* \ll K_{ij}$. Under this regime, we can approximate the growth rate to be simply proportional to resource concentration:

$$\frac{1}{N_i}\frac{dN_i}{dt} = \sum_j r_{ij}\, \frac{c_j}{K_{ij} + c_j} - \delta \simeq \sum_j \frac{r_{ij}}{K_{ij}} c_j - \delta \qquad Eqn. 21$$

Here, the equilibrium resource concentration becomes:

$$c_{ij}^* = \delta \frac{K_{ij}}{r_{ij}} \qquad Eqn. 22$$

The condition for species 1 to win in resource 1 while species 2 wins in resource 2 is then:

$$\frac{K_{11}}{r_{11}} < \frac{K_{21}}{r_{21}}, \qquad \frac{K_{12}}{r_{12}} > \frac{K_{22}}{r_{22}} \qquad Eqn. 23$$

***Condition 2***: High dilution regime: $\delta < r_{ij},\ \delta \sim r_{ij}$

In this case, $\delta$ is large and approaches (but does not exceed) the maximum growth rate for each species on each resource. This limit is necessary in order to obtain positive solutions for $c_{ij}^*$:

$$c_{ij}^* = \frac{\delta K_{ij}}{r_{ij} - \delta} \qquad Eqn. 24$$

Then the condition for niche flip becomes:

$$\frac{K_{11}}{r_{11} - \delta} > \frac{K_{21}}{r_{21} - \delta}, \qquad \frac{K_{12}}{r_{12} - \delta} < \frac{K_{22}}{r_{22} - \delta} \qquad Eqn. 25$$

Focusing on resource 1, combining the condition for low- and high-disturbance leads to:

$$\frac{r_{11}}{r_{21}} > \frac{K_{11}}{K_{21}} > \frac{r_{11} - \delta}{r_{21} - \delta} \qquad Eqn. 26$$

Under the assumption that $r_{ij} > \delta$, these inequalities are equivalent to:

$$r_{21} > r_{11},\ K_{21} > K_{11}\frac{r_{21}}{r_{11}} \qquad Eqn. 27$$

Recall that in the high-dilution regime, species 2 wins in resource 1 and species 1 wins in resource 2. Similarly, for resource 2, we have another inequality in the reverse direction:

$$r_{22} < r_{12},\ K_{22} < K_{12}\frac{r_{22}}{r_{12}} \qquad Eqn. 28$$

Thus, for dilution regimes where each resource is capable of supporting each species' survival, two requirements must be met in order to enable niche flip. First, at a given dilution rate, each species should win on a different resource. Second, the outcome of competition on each resource should flip between low and high dilution regime, which is possible if *r*/*K* tradeoffs occur on each resource.

Additionally, based on these results, we can build intuition about multi-species, multi-resource communities, similar to those in our experiments. In order to exclude all instances of niche flip that would lead to a U-shaped DDR in a multi-species community in complex media, every pair of species must not exhibit *r*/*K* tradeoffs in all pairs of resources. This implies that in order for the community to not exhibit a U-shaped DDR, the tradeoff condition (Eqn. 25 above) should not be met among all pairs in all resources, and instead the following reverse condition should be satisfied for all $i, j, k$:

$$r_{ij} < r_{kj}\ \ iff\ \frac{r_{ij}}{K_{ij}} < \frac{r_{kj}}{K_{kj}} \qquad Eqn. 29$$

Furthermore, we can extend this condition to overall parameters measured in a complex media, rather than on each component resource. Summed Monod growth on all resources can be approximated by a single Monod growth curve with:

$$r_i = \sum_j r_{ij}\,, \qquad K_i = \frac{r_i}{\sum_j \frac{r_{ij}}{K_{ij}}} \qquad Eqn. 30$$

To satisfy the condition for avoiding a U-shaped DDR, the community should show a positive correlation between $r_i$ and $r_i/K_i$:

$$r_i < r_k\ \ iff\ \frac{r_i}{K_i} < \frac{r_k}{K_k} \qquad Eqn. 31$$

In conclusion, in the summed Monod model, a U-shaped DDR is guaranteed when $r$ and $r/K$ in complex media do not have a positive correlation. To be certain, the strength of the U-shape should depend on the prevalence of tradeoffs, and by extension, a weak correlation would result in a less dramatic U-shape than would a strong negative correlation. Additionally, observing a U-shape resulting from a small number of rare tradeoffs would require measuring the community at the dilution rates where the few instances of niche flip occur. The negative correlation exhibited by our isolates (Supplementary Fig. 13), however, indicates that niche flip is a relevant mechanism that warrants further investigation.

Thus far, we derived the required conditions for niche flip in a two-species, two-resource community:

$$r_{21} > r_{11},\ K_{21} > K_{11}\frac{r_{21}}{r_{11}},\ r_{22} < r_{12},\ K_{22} < K_{12}\frac{r_{22}}{r_{12}} \qquad Eqn. 32$$

These inequalities represent trade-offs between $r$ and $K$ that cause the Monod growth curves of the two species in both resources to cross, but such that the species that grows faster at low resource concentration flips between the two resources (Supplementary Fig. 12).

As discussed in the main text, these $r/K$ tradeoffs make coexistence possible at low and high disturbances, while at some level of intermediate disturbance the niche flip eliminates the possibility for coexistence. This mechanism could lead to a U-shaped diversity-disturbance relationship among a community of many species consuming many resources, assuming that trade-offs between $r$ and $K$ were prevalent. In this section, we show that this requirement can be overcome, as $r/K$ trade-offs across different resources can also lead to niche flip. To illustrate this concept, we must consider an even higher disturbance regime than explored previously.

Let us first check whether eliminating $r/K$ tradeoffs on each resource would eliminate the U-shaped DDR. In a simulation of a 10-species, 7-resource community, we sorted $r$ and $K$ such that the Monod growth curves on each of the seven resources do no intersect, in order to avoid trade-offs in each of the seven resources. An example of the Monod curves on one of the resources is shown below:

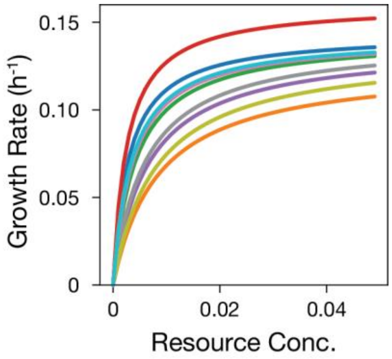


**Supplementary Figure 5. Generating species with sorted $r$ and $K$ parameters prevents intersection of growth curves.** Growth curves of 10 species on a single resource are shown. For these species in a single resource environment, the outcome of competition does not depend on resource concentration or dilution rate.

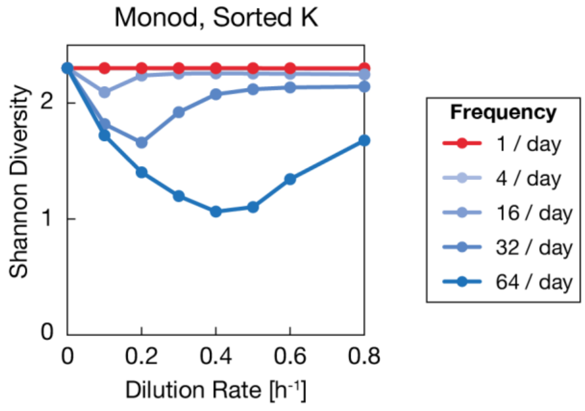


**Supplementary Figure 6. Communities with sorted $K$ per resource still exhibit U-shaped DDR.** Shannon diversity of communities vs. mean dilution rate. Color indicates frequency of fluctuations. Calculations performed after 6 days of simulated growth, and averaged across 100 communities with randomly generated and sorted $r/K$ parameters.

Surprisingly, this sorting did not eliminate the overall U-shaped DDR. To explain how a lack of $r/K$ tradeoffs on each resource could nevertheless lead to niche flip and the U shape, we return to the example of a two-species, two-resource community. Let us consider an even higher disturbance regime than explored previously.

***Condition 3***: Very high dilution regime: $\delta > r_{ij}$

In this regime, the dilution rate is greater than any species' maximum growth rate on any single resource. In the summed Monod model, species can survive when the sum of growth rates across resources can overcome the dilution rate.

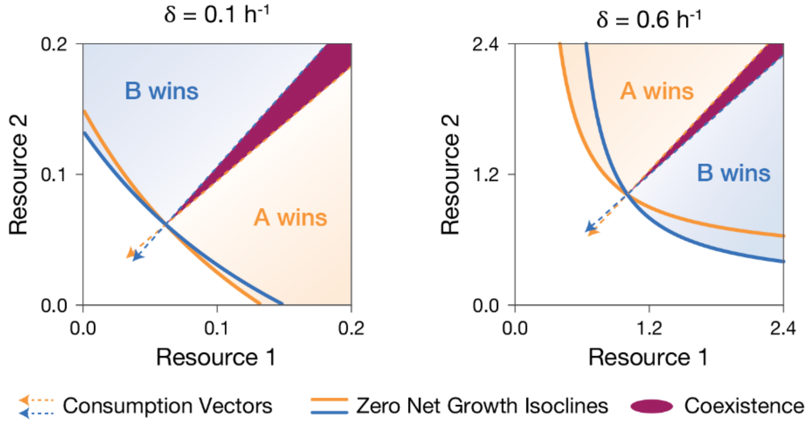


**Supplementary Figure 7. Niche flip occurs in a community with sorted $K$ parameters at high dilution rate.** At low dilution rate, ZNGIs intersect the axes and the resource consumption vectors outline a coexistence region. At high dilution rate, species cannot survive on a single resource, so ZNGIs do not intersect the axis. The relative positions of ZNGIs and consumption vectors (i.e. invasion boundaries) have flipped at high dilution rate, consistent with niche-flip.

In the two-species, two-resource example, neither species can survive on a single resource, but both could survive in the presence of both resources at a sufficient concentration. The niche of each species is now determined not by the intersection of the ZNGIs and the resource axis, but by the asymptotic distance between the ZNGI and the resource axis. A species that requires less additional resource 2 in order to survive would win in media containing mostly resource 1. For this reason, growth on a second resource now plays a role in determining the winner in the (predominantly) first resource. Under the summed Monod model, the necessary concentration of resource 2 can be solved explicitly:

$$r_{i1} + r_{i2}\frac{c_{i2}^*}{K_{i2} + c_{i2}^*} = \delta \qquad Eqn.33$$

Solving this gives:

$$c_{i2}^* = K_{i2}\ \frac{\delta - r_{i1}}{r_{i1} + r_{i2} - \delta} \qquad Eqn.34$$

As a reminder, in the low dilution limit, the competition outcome on resource 1 is determined by:

$$c_{i1}^* = K_{i1}\frac{\delta}{r_{i1}} \qquad Eqn.35$$

Without losing generality, let us assume species 1 outcompetes species 2 at low dilution rates in resource 1. Then the niche flip requires that the outcome flips as we change the dilution rate and species 2 outcompetes species 1 in resource 1 under the high-dilution regime. Then the niche flip condition becomes:

$$\frac{K_{11}}{r_{11}} < \frac{K_{21}}{r_{21}}\ , \qquad K_{12}\frac{\delta - r_{11}}{r_{11} + r_{12} - \delta} > K_{22}\frac{\delta - r_{21}}{r_{21} + r_{22} - \delta} \qquad Eqn.36$$

Now the competition outcome under large disturbance depends not only on the maximum growth rate on the particular resource but also on the Monod constant on the other resource, which determines the extra amount of the other resource necessary to overcome the dilution rate. This coupling between two resources opens up additional conditions for niche flip without $r/K$ trade-offs in each resource. For example, under simplifying assumption that $r_{11} + r_{12} = r_{21} + r_{22}$, one can have niche flip with

$$r_{11} > r_{21},\ \ K_{12} < \frac{\delta - r_{12}}{\delta - r_{22}}K_{22}, \qquad r_{12} < r_{22},\ \ K_{11} > \frac{\delta - r_{11}}{\delta - r_{21}}K_{21} \qquad Eqn.37$$

In this example, there are no apparent $r/K$ trade-offs on each resource. The trade-off between $r_{i1}$ and $K_{i2}$ leads to the flipped outcome on resource 1, and similarly the trade-off between $r_{i2}$ and $K_{i1}$ leads to the flipped outcome on resource 2.

Therefore, since $r/K$ trade-offs on separate resources can also lead to niche flip, we must keep $K$ constant for all species grown on any particular resource in order to eliminate the possibility of niche flip. In a simulation of a ten-species, seven-resource community, with one $K$ per resource assigned to all ten species, the diversity-disturbance relationship is not U-shaped:

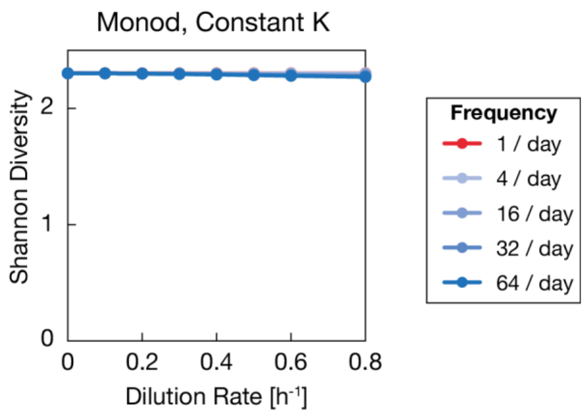


**Supplementary Figure 8. Communities with constant $K$ across all resources do not exhibit U-shaped DDR.** Shannon diversity of communities vs. mean dilution rate. Color indicates frequency of fluctuations. Calculations performed after 6 days of simulated growth, and averaged across 100 communities with randomly generated maximal growth rate $\boldsymbol{r}$, and a randomly generated $\boldsymbol{K}$, shared by all species.

In conclusion, we observe that when the dilution rate is smaller than the maximal growth rate on each resource, $r/K$ tradeoffs on each resource are required for niche-flip and a U-shaped diversity-disturbance relationship. If we consider an even higher dilution rate, $r/K$ tradeoffs on each resource are no longer required for niche-flip because $r/K$ tradeoffs across different resources can also lead to niche-flip. This opens up additional parameter space that would lead to a U-shape diversity-disturbance relationship.

In a multi-resource multi-species community, this implies that even a positive correlation between $r$ and $r/K$ in complex media does not necessarily guarantee the absence of a U-shaped DDR. Our results show that niche flip and a resulting U-shaped DDR under constant disturbance are exceedingly difficult to avoid in the Monod model, and that their absence requires a very constrained parameter space. Note however, that in this section we have considered only chemostat continuous dynamics. As shown experimentally and in simulations, introducing fluctuations into the system entirely reshapes the DDRs (**Fig. 3e and 3f**). Finally, while the simulations are robust to a variety of parameter choices and alternative formulations (Supplementary Figures 3 and 4), populating the model with growth parameters from different species-media combinations may yield a diverse range of DDR shapes. Nevertheless, we believe the interesting features of the Niche-flip mechanism could warrant its study in other ecosystems.

### Supplementary Discussion 3: Additional Experimental Results

DDR64 Experiment

As discussed in the main text, the DDR64 Experiment showed that disturbance intensity and fluctuations have distinct effects on both composition and diversity when applied independently. Additionally, this experiment highlights the reproducibility of ecosystems grown in controlled conditions of continuous culture, as we observed a high degree of agreement between replicates. There are several additional factors to consider that were not covered in entirety in the main text.

Though we focused on Shannon diversity from 16S sequencing, we found that the general results were consistent between diversity metrics: Shannon diversity of 16S ASVs, richness of 16S ASVs, and richness of colony morphotypes (Supplementary Fig. 17). While counting colony morphotypes cannot achieve the measurement depth of sequencing, nor does it provide complete composition measurement (*Chromobacterium*, though abundant in 16S measurements were not isolated on our plates, for example), it is encouraging to see agreement.

Though not the focus of this study, the dynamics of both composition and diversity are of interest. We collected samples at daily timepoints, and therefore were able to track changes in composition over time (Supplementary Figs. 14 and 15) with corresponding changes in diversity (Supplementary Fig.16). As expected, the community undergoes significant selection for growth in Nutrient Broth in eVOLVER, as shown by comparison to the compositions of soil, the PBS immersion, and the post-enrichment samples used to inoculate the DDR64 Experiment (Supplementary Fig. 10). We also note the putative succession occurring in the DDR64 Experiment, as many cultures exhibit transient similarity in composition at the Day 1 or Day 2 timepoints (Supplementary Figs. 14 and 15). These compositions are distinct from both the initial inoculum and the endpoint compositions. This transient community is also correlated with transient increases in Shannon diversity in many conditions (Supplementary Fig. 16). Finally, we note that while it is not possible to determine whether the endpoint composition is representative of true equilibrium, changes in composition and diversity do appear to slow as time goes on, particularly by days 4-6. It is notable that while communities grown at different dilution rates must undergo different numbers of generations in order to survive the same number of days, all communities seem to approach this quasi-steady state at similar rates. The Monod consumer resource model also predicts similar qualitative DDRs across a wide range of simulation lengths (Supplementary Fig. 3), so we feel the experimental results are informative, not merely transient.

We note that while the Monod consumer resource model can explain our results, there are potential confounding factors that could be at work. Firstly, we measured differences in biofilm accumulation across the parameter space using optical density measurements in eVOLVER as a crude metric (Supplementary Fig. 18). After flushing the vial with fresh media, non-zero optical density measurements can be attributed to biofilm on the walls of the glass culture vial. By attaching to the wall, a species could escape death by dilution and therefore skew composition and diversity. Accordingly, we did observe discrepancies between OD, CFU/mL and DNA extractions that may be partially correlated with biofilm accumulation (Supplementary Fig. 20). However, we did not observe biofilm accumulation to be correlated with diversity (Supplementary Fig. 19), indicating that differences in biofilm are correlative rather than causative. In future experiments, the use of surfactants (Tween) could be used to mitigate this effect, though these may have other unintended effects on the community. Similarly, changes in pH, oxygenation, phage, or antimicrobial levels could conceivably be linked to diversity changes and were not measured in this experiment. Nevertheless, we are encouraged that the Monod consumer resource model qualitatively explains our results without invoking any of these factors. Notably, modifying the model to include asymmetric mortality rates did not qualitatively change the shape of DDRs in simulations (Supplementary Fig. 4b), suggesting that our results could be robust to these types of effects. As discussed in the main text, if the growth response of each species to these types of factors can be mapped to dilution rate in a Type II functional response, these effects may interact with disturbance intensity and fluctuations in a manner similar to the consumer resource models we have focused on.

DDR Washout Experiment

After observing large differences across the parameter space in the DDR64 Experiment, we wanted to run a similar experiment to answer two questions: 1) are the compositions we observed reproducible, or idiosyncratic, and 2) what happens at more extreme levels of disturbance? As discussed in the main text, by repeating the 0.3 $h^{-1}$ dilution rate condition, we indeed found that compositions and diversity were reproducible, and that diversity did increase at higher dilution rates, consistent with the U-shaped DDR. The second question was harder to answer.

Intuition and simulations (**Fig. 4**) predict that when dilution rates far exceed the growth rates that can be achieved by species in the ecosystem, these species will be diluted out over time. At sufficiently high disturbance rates, no community member survives (termed "washout"), and diversity drops to zero. Similarly, for fluctuating conditions, disturbances of sufficient size could bring population levels low enough that compositions are dominated by stochastic bottleneck events rather than deterministic competition. In our 48-vial DDR Washout experiment, we ran dilution rates as high as 1.5$h^{-1}$, higher than the measured growth rate of most isolates from the experiment (Supplementary Fig. 13). In eVOLVER, this represents the technical limit of dilution rates for a 1/day disturbance (running pumps approximately 50% of the time for a safety factor). Despite the dilutions theoretically washing out most species and imposing dilution factors larger than the population size in each vial, we did not observe complete washout (Supplementary Fig. 11). As discussed earlier, biofilm represents one possible way that species could escape washout. Also, though eVOLVER vials are constantly stirred and evenly mixed at low dilution rates, when dilutions are very fast, the mixing timescale may be slow in comparison. This breaks the well-mixed assumption, so cells could escape dilution. Therefore, we were cautious to draw strong conclusions from this part of the parameter space.

Growth Characterization Experiments

The DDR64 and DDR Washout experiments were performed from an undefined starting composition, in dilute Nutrient Broth, a complex media with diverse nutrient sources (e.g. amino acids, peptides, nucleic acids). While natural communities are often similarly composed of many species and nutrients, this design makes it very challenging to identify the specific species-nutrient combinations with $r/K$ tradeoffs capable of generating the observed DDR. The most robust test of the niche-flip mechanism would be to conduct co-culture experiments with species that have measured growth parameters in defined media, then compare results to simulated competition outcomes. Such an experiment first requires accurate measurement of Monod growth parameters ($r$ and $K$). We set out to measure these values for isolates from the DDR64 Experiment. Specifically, if we found $r/K$ tradeoffs that varied between resources, these could be used to support and/or directly test the niche-flip mechanism.

We first asked whether isolates exhibited saturating growth in the media composition used in our experiment. In 96-well plates, we measured growth rates at varying concentrations of diluted Nutrient Broth. Growth rates varied as a function of nutrient composition, and we found that growth rates indeed saturated at nutrient levels used in the experiment (Supplementary Fig. 13). Fitting Monod parameters to these measurements, we also observed $r/K$ tradeoffs in isolates grown on nutrient broth (Supplementary Fig. 13). However, it is unclear what the limiting nutrient is in the Nutrient Broth is, as the complex media is comprised of peptone and yeast extract, and therefore we sought to repeat these experiments in defined minimal media, where we could titrate a single limiting nutrient.

We repeated the above experiment in carbon-limited M9 minimal media supplemented with varying concentrations of amino acids as a sole carbon source. We observed lower levels of growth in M9, and growth at lower concentrations of amino acid was below the limit of detection of our microplate spectrophotometer, preventing us from fitting Monod growth parameters. Nevertheless, we observed that at saturating carbon concentrations, the isolate with higher maximal growth rate varied for different amino acids (Supplementary Fig. 13). This is consistent with $r/K$ tradeoffs, though it is not sufficient evidence on its own.

Regardless, these two experimental observations support our hypothesis that the species and conditions used in our experiments exhibit Monod growth kinetics. This means that 1) DDRs would be expected to be reshaped by fluctuations, and 2) species could feasibly undergo niche-flip to yield the U-shaped DDRs under constant disturbance.

To enable the types of direct tests described at the beginning of this section, more characterization work is needed. Specifically, methods for growth measurement that retain sensitivity at low resource concentrations are needed. Furthermore, these methods must be of sufficient throughput to enable measurement across multiple species, nutrients, and nutrient concentrations. Advances in microfluidics, specifically droplet microfluidics may soon enable these types of studies[54,55]. In the absence of these experiments, we are still encouraged that our measurements do not invalidate the niche-flip mechanism and in fact support it. Additionally, as discussed in Supplementary Discussion 3, U-shaped DDRs are possible in the Monod-growth model even without the $r/K$ tradeoffs we sought in these experiments, which further increases the feasibility of the model.

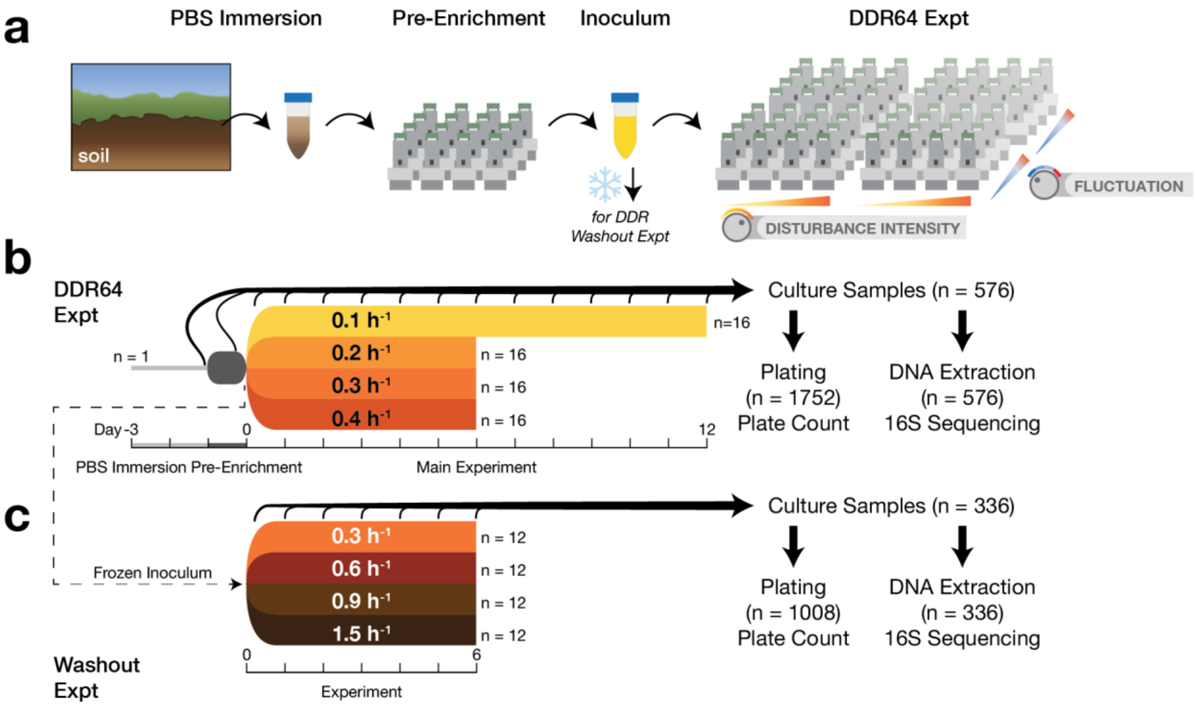


**Supplementary Figure 9. Timeline of DDR64 and DDR Washout Experiments. (a)** Soil samples were immersed in PBS, then pre-enriched in overnight cultures in 0.1X Nutrient Broth. Post-enrichment cultures were mixed to form the final inoculum. The final inoculum was also frozen for follow up experiments. **(b)** Timeline for the DDR64 Experiment. Cultures were sampled daily for plating and sequencing to determine composition. Cultures completed ~20-90 generations over 6 days. The 0.1 $h^{-1}$ cultures were carried out for an additional 6 days in order to complete an additional 20 generations, for a total of 40. **(c)** Timeline and sampling schedule for DDR Washout Experiment. This experiment was started from a frozen sample of the inoculum used in the DDR64 experiment.

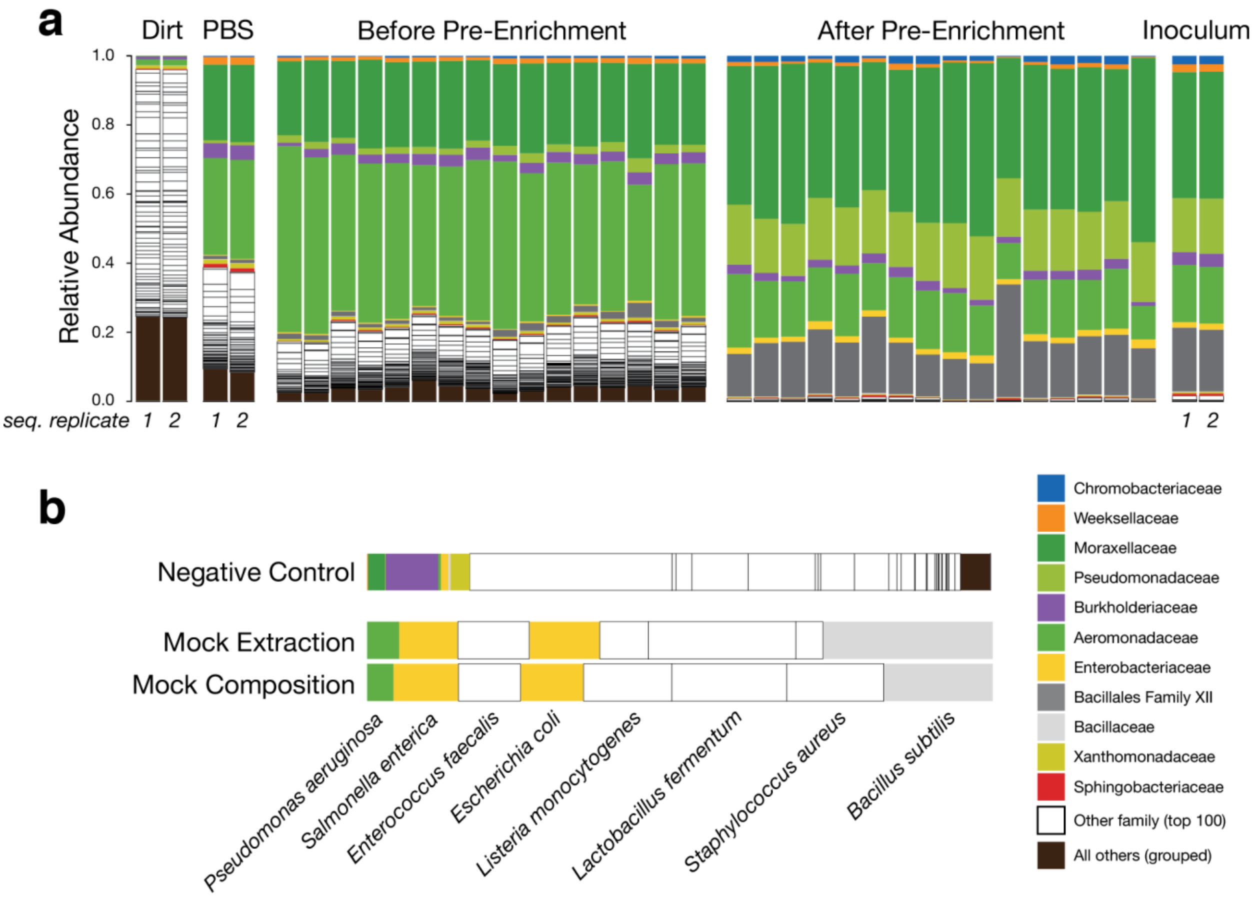


**Supplementary Figure 10. Inoculum composition determined by 16S sequencing. (a)** Family-level composition of a representative soil sample, supernatant of PBS immersion, pre-enrichment cultures, and final inoculum. Diversity and composition changes over time indicate selection for growth in lab conditions. **(b)** Composition of negative control sample, positive control mock community sample, and the true composition of the mock community sample. Positive control samples indicate coverage over both Gram-positive and Gram-negative organisms. Negative control sample indicates that the composition of experimental samples is significantly different, as would be expected for samples of sufficient biomass.

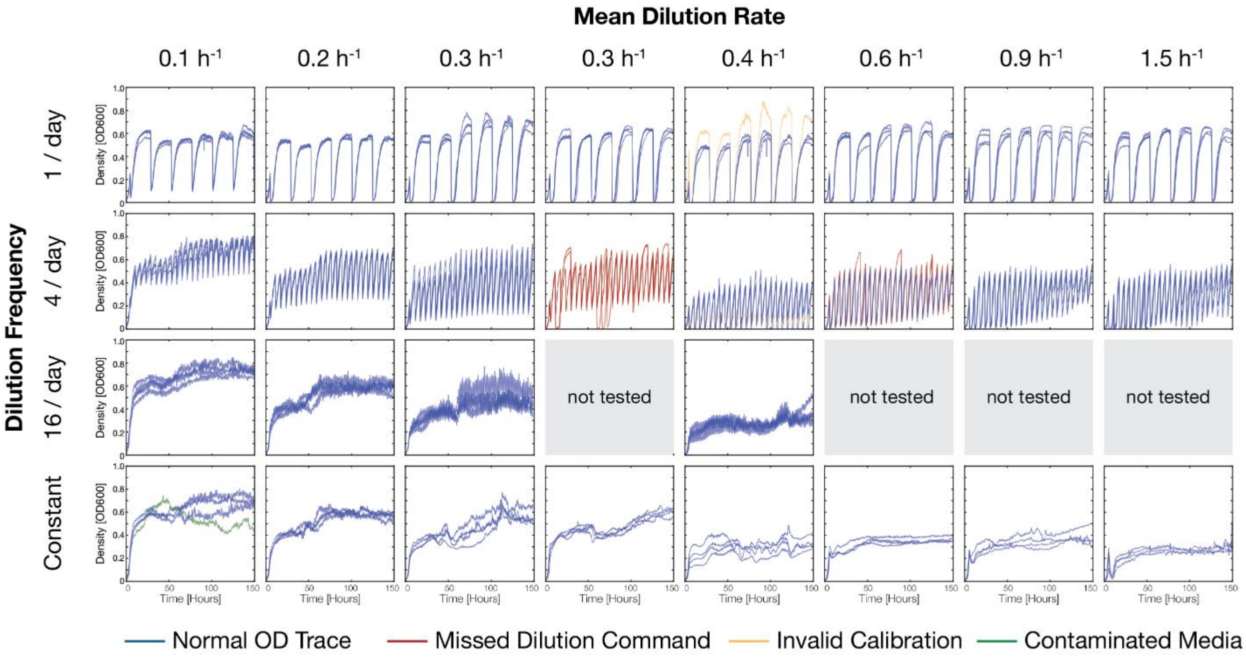


**Supplementary Figure 11. Optical density over time for DDR64 and DDR Washout experiments.** Optical density was recorded in each eVOLVER vial over time at ~30 second intervals. Plots from vials in replicate conditions are included on the same axes. Colors used to indicate technical issues in that vial. DDR64 experiment located in columns 1,2,3,5; DDR Washout experiment in columns 4,6,7,8.

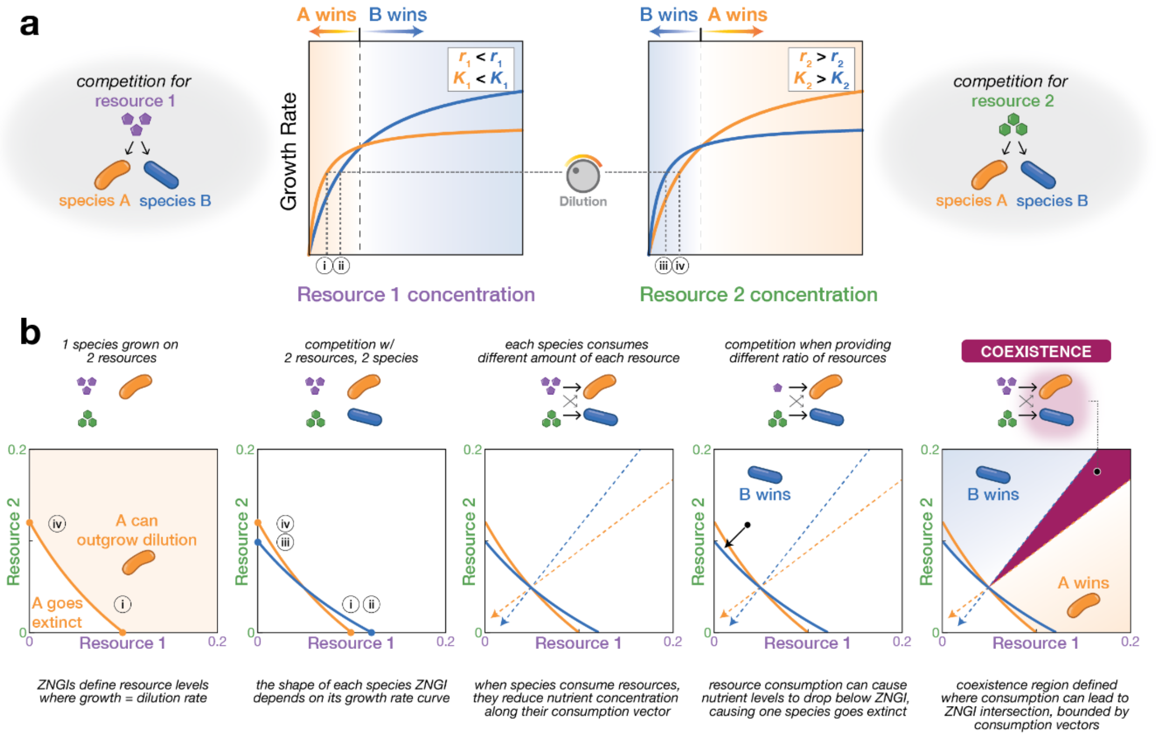


**Supplementary Figure 12. Navigating between Monod growth curves and Tilman diagrams.** Growth rates calculated using consumer resource models can be used to construct diagrams of competitive outcomes, see Tilman[37]. **(a)** Consider Species A and Species B, which each exhibit different resource concentration-dependent growth on Resource 1 or Resource 2. To achieve growth rates equal to this indicated dilution rate, the species require different resource concentrations to survive (labelled i-iv). **(b)** In a multi-resource environment, the summed growth rate across resources constructs a Zero Net Growth Isocline (ZNGI) which intersects the axes at points (i-iv) from the prior panel. ZNGIs define the space of supplied resource that can support a species sufficiently such that its growth meets or exceeds the mortality rate (e.g. dilution rate). The outcome of competition can be predicted by determining whether resource consumption will drive one species to extinction or not. The borders of these different outcomes, termed invasion boundaries in Figure 2, can be identified by calculating a resource consumption vector at the intersection of the ZNGIs. Note that Tilman diagrams may be constructed for any consumer resource model, and the shapes and movement of the ZNGIs will change accordingly.

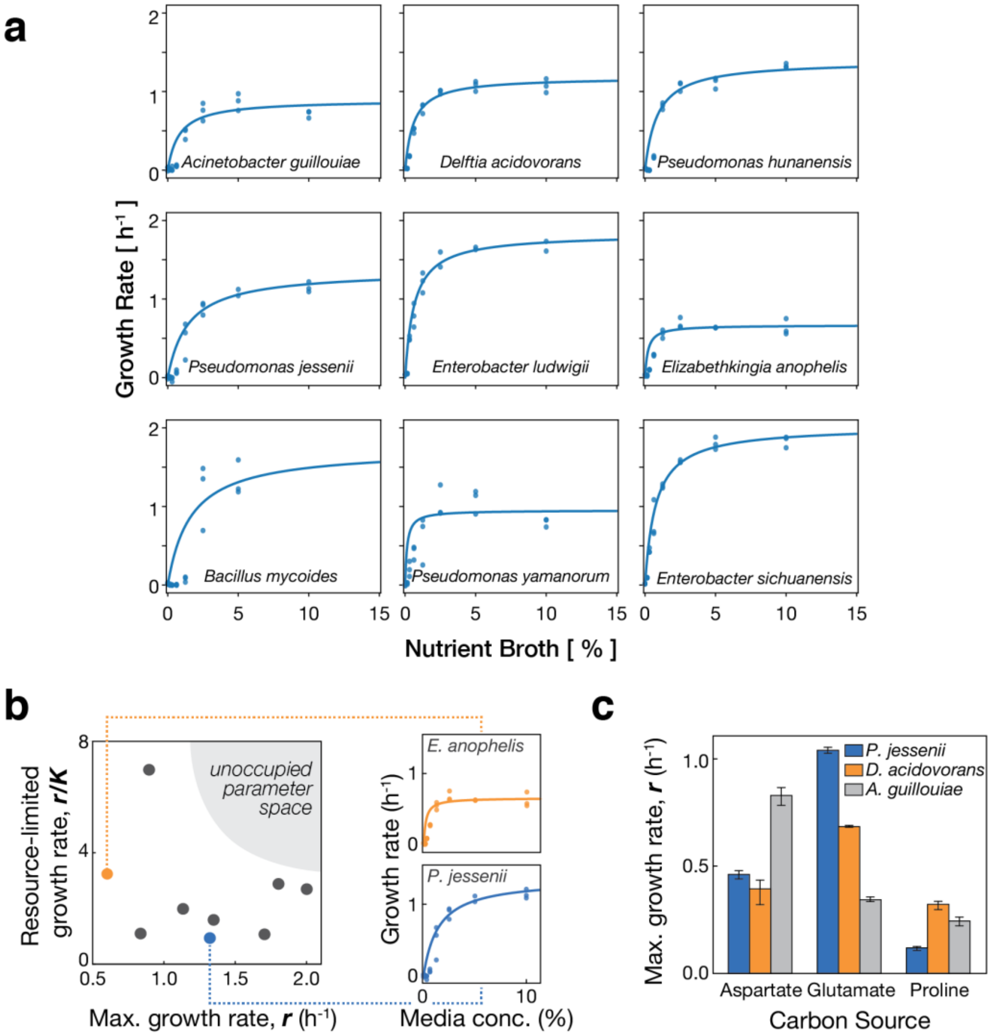


**Supplementary Figure 13. Measured growth of isolates in Nutrient Broth indicate variation in r and K parameters across different medias. a)** 9 isolates from the DDR64 and DDR Washout experiments were grown in a microplate spectrophotometer in dilute Nutrient Broth media (ranging from 0.1X to <0.0016X, plus a water-only negative control). Linear regression on a Lineweaver-Burk plot was used to calculate Monod $\boldsymbol{r}$ and $\boldsymbol{K}$ values. Monod curves generated from those fit values are indicated by solid lines, while triplicate growth measurements are indicated as points. Flattening of growth rate vs. nutrient concentration curves is consistent with saturating growth in Monod model. **b)** Scatter plot of Monod $\boldsymbol{r}$ and $\boldsymbol{r/K}$ values indicate that while species vary in Monod $\boldsymbol{r}$ and $\boldsymbol{K}$, the unoccupied parameter space indicates that no species would be expected to outcompete in both resource rich and resource poor environments. **c)** 3 isolates from the DDR64 and DDR Washout experiments were grown in M9 media supplemented with 10mM of the indicated amino acid as a sole carbon source. Different isolates had different growth rates on each resource, indicating resource preferences.

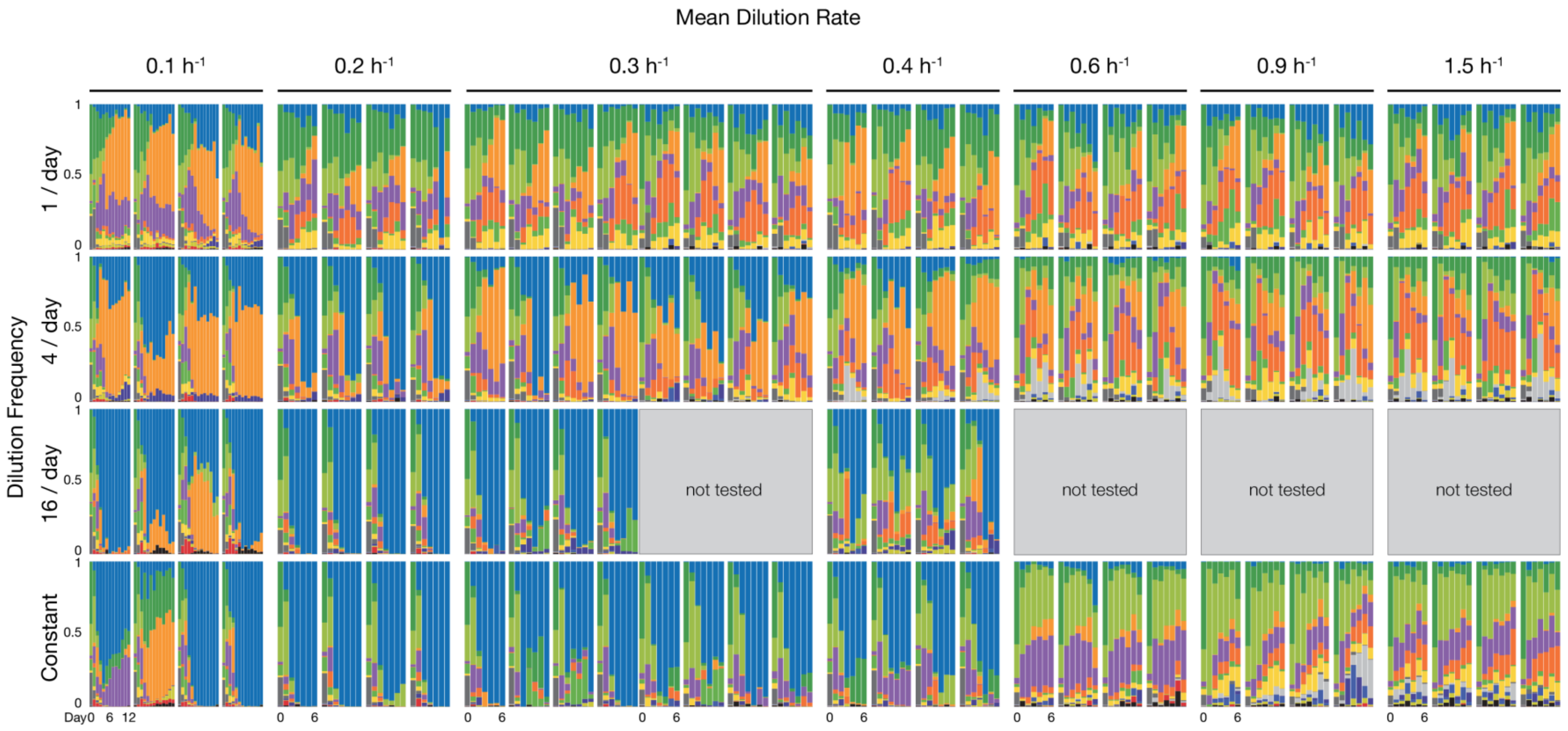


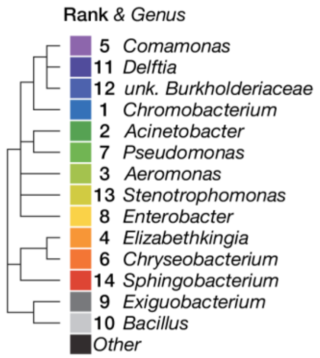


**Supplementary Figure 14. Genus-level composition over time in the DDR64 and DDR Washout experiments.** Composition was determined to vary over time across different disturbance intensity and fluctuation levels. Daily samples were taken from soil microbe co-cultures grown in eVOLVER in varying dilution profiles (Materials and Methods). Amplicon sequencing on the v4 16S region was performed and analyzed in QIIME 2 to assign genus level taxonomy according to the SILVA 132 database. Each bar depicts the relative abundance of genera in a single sample. Mean rank abundance of a bacterial genus (across all samples) is denoted by order (top to bottom), and taxonomic similarity is denoted by color (see legend).

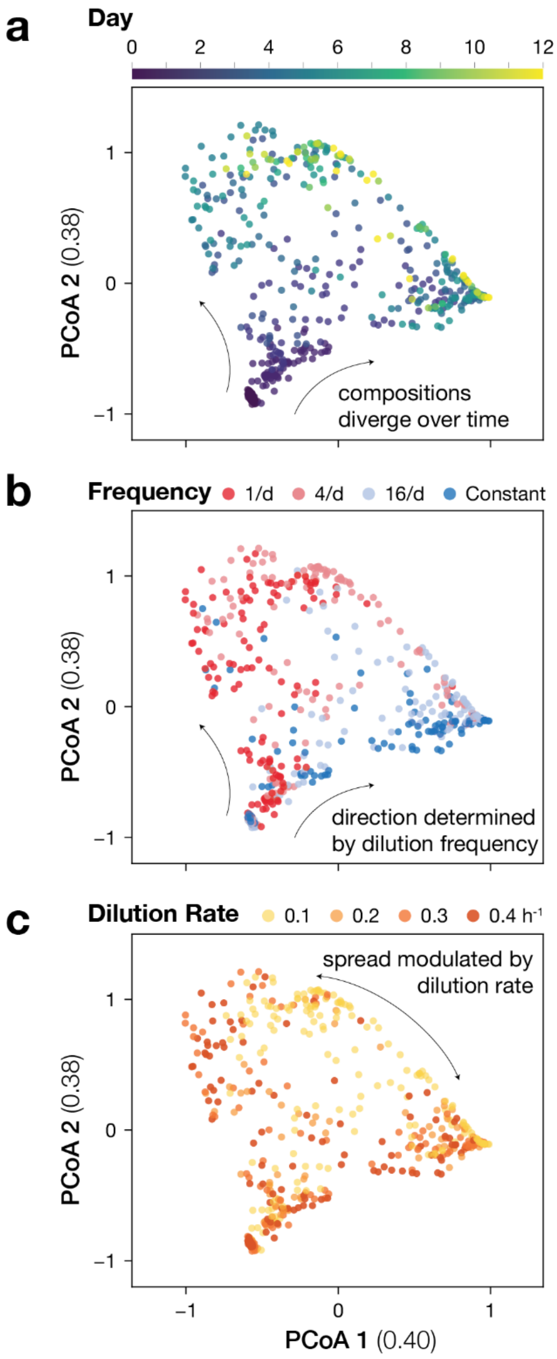


**Supplementary Figure 15. Principle Coordinate Analysis shows varying composition in DDR64 Experiment.** Principal Coordinate Analysis indicates divergence in composition between samples. Ordination was based on Weighted UniFrac distances that were calculated from 16S composition of daily samples in QIIME 2. **(a)** Samples colored according to day of collection. **(b)** Samples colored to indicate the dilution frequency of the culture. **(c)** Samples colored to indicate the mean dilution rate of the culture. The magnitude for each Principle Coordinate is listed in parentheses.

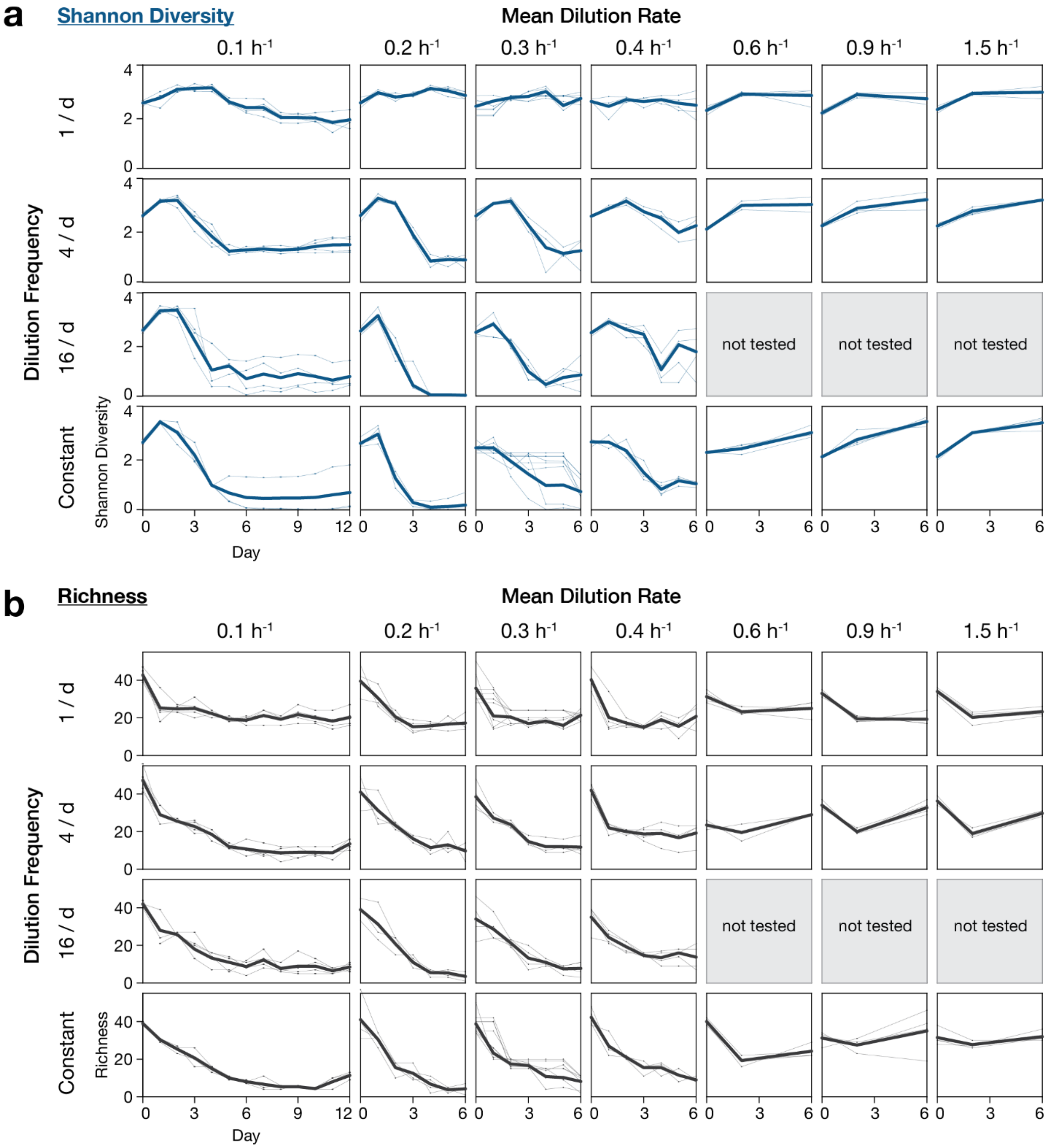


**Supplementary Figure 16. Diversity changes over time in DDR64 and DDR Washout experiments.** Diversity was calculated from 16S sequencing for each timepoint sample in each vial. **(a)** The Shannon diversity of 16S ASVs for each vial are indicated by points connected by thin lines. The mean across replicate vials is denoted with a thick line. **(b)** 16S ASV richness for each vial, plotted as above. Samples with technical issues (e.g. poor sequencing coverage, contamination, see Supplementary Fig. 11) were excluded from analysis.

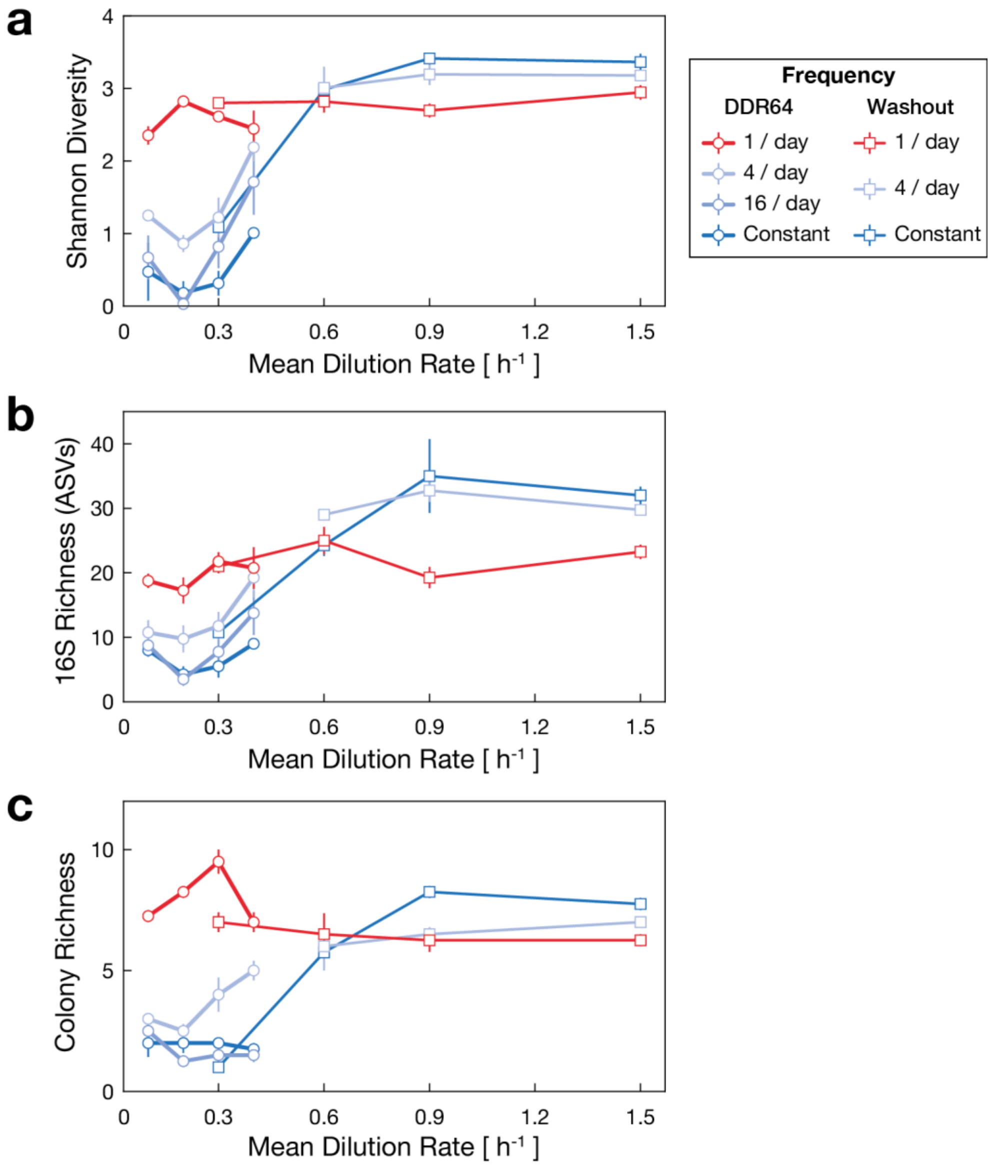


**Supplementary Figure 17. Different diversity metrics from DDR64 and DDR Washout experiments show U-shaped DDRs.** Diversity was calculated using the Day 6 timepoint for all vials in DDR64 and DDR Washout experiments. **(a)** Shannon diversity of 16S ASVs for each vial, mean across replicates. **(b)** 16S ASV richness for each vial, mean across replicates. **(c)** Colony richness, the number of unique colony morphotypes identified on Nutrient Agar plate for each vial, mean across replicates. Note that due to CFU/mL differences, colony richness is deflated in low density conditions. This effect is pronounced in the constant dilution vials. Bars indicate standard error of the mean. Samples with technical issues (e.g. poor sequencing coverage, contamination, see Supplementary Fig. 11) were excluded from analysis.

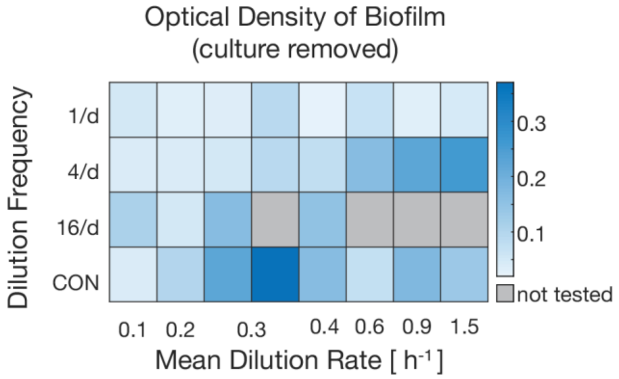


**Supplementary Figure 18. Heatmap of biofilm levels at endpoint.** Biofilm levels were measured by flushing vials with fresh media then measuring optical density in eVOLVER. Vials with higher optical density have more biofilm accumulation. Heatmap depicts mean across replicates. DDR64 experiment data in columns 1,2,3,5 and DDR Washout experiment in columns 4,6,7,8.

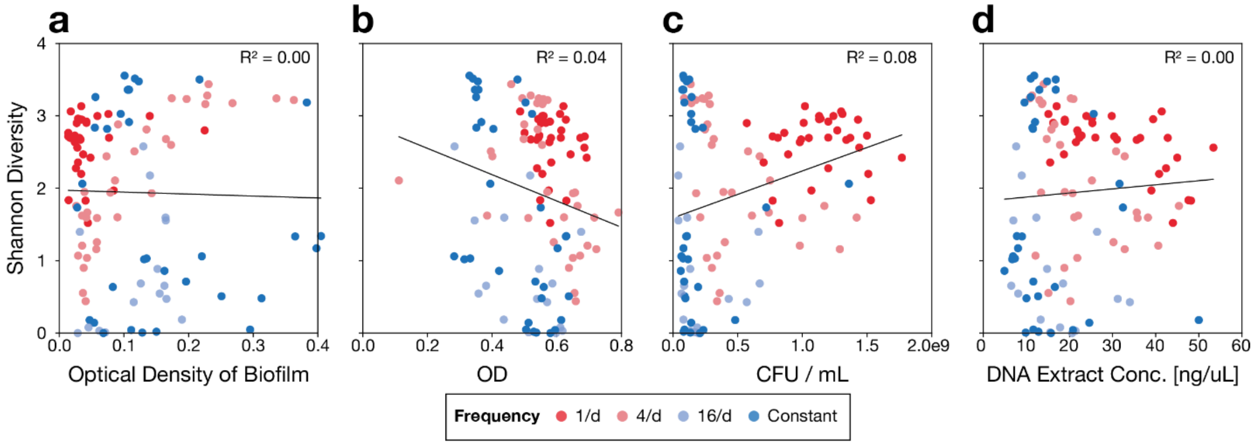


**Supplementary Figure 19. Diversity does not correlate with potential confounding factors.** Shannon diversity of 16S ASVs for endpoint samples plotted against potential confounding factors. **(a)** Diversity vs. Optical Density of biofilm (see fig. S14). **(b)** Diversity vs. Optical Density in eVOLVER. **(c)** Diversity vs. CFU/mL calculated from Nutrient Agar plate. **(d)** Diversity vs. concentration yield of DNA extraction. For all plots, a linear regression line is plotted in black, with corresponding $R^2$ values. Color indicates frequency of dilution (see legend).

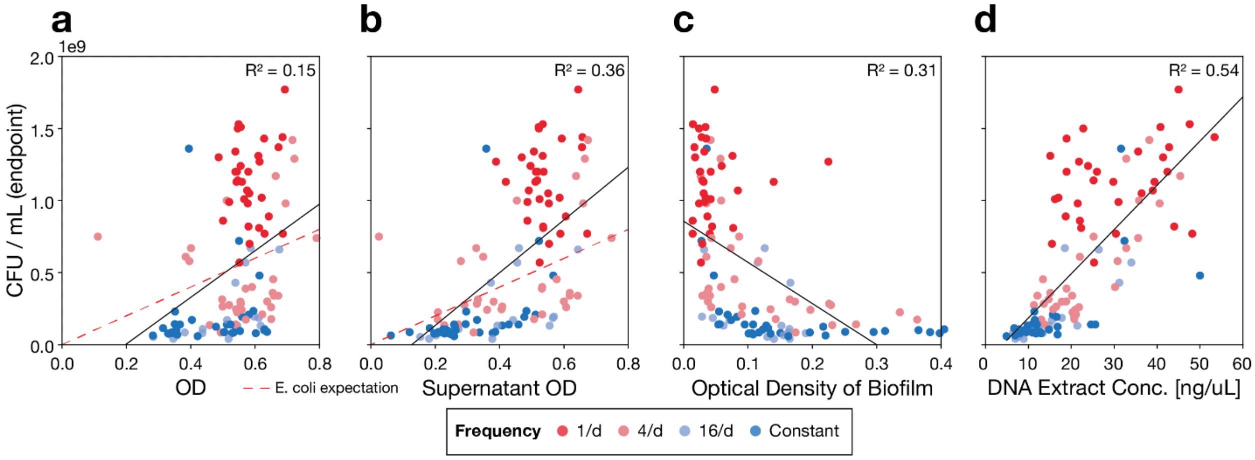


**Supplementary Figure 20. Correlation between population size estimates vary according to condition and biofilm accumulation.** Colony forming units (CFU) for endpoint samples plotted against other population size estimates. **(a)** CFU/mL vs. Optical Density in eVOLVER. **(b)** CFU/mL vs. Optical Density in eVOLVER, minus biofilm contribution. **(c)** CFU/mL vs. Optical Density of biofilm (see fig. S13). **(d)** CFU/mL vs. concentration yield of DNA extraction. For all plots, linear regression line is plotted in black, with corresponding $R^2$ values. For optical density plots, a linear relationship (CFU/OD = 8 x $10^8$) Color indicates frequency of dilution (see legend). Samples in the 1/d condition group were measured to have much higher CFU/mL, as these cultures reached saturation.
